# Integrative transcriptome-based drug repurposing in tuberculosis

**DOI:** 10.1101/2025.06.02.657296

**Authors:** Kewalin Samart, Ling Thang, Landon Buskirk, Amy Tonielli, Arjun Krishnan, Janani Ravi

## Abstract

Tuberculosis (TB) remains the leading cause of infectious disease mortality worldwide, killing over one million people annually. Rising antibiotic resistance has added urgency to the need for host-directed therapeutics (HDTs) that modulate host immune responses alongside directly targeting the pathogen. Repurposing FDA-approved drugs is particularly attractive for this purpose because their safety profiles are already well established, substantially reducing development time and cost. Transcriptomic methods have successfully identified repurposable therapeutics for TB based on ‘connectivity mapping,’ which identifies drugs that reverse disease gene expression patterns. However, these applications are limited to a small subset of data belonging to a specific data platform and a few connectivity methods. Expanding beyond these constrained settings introduces substantial challenges, including dataset heterogeneity across transcriptomics platforms and biological conditions, uncertainty about optimal scoring methods, and the lack of systematic approaches to identify robust disease signatures. We developed a computational workflow that integrates 28 TB gene expression signatures and multiple connectivity scoring methods to capture dominant TB signals regardless of variation in microarray and RNAseq platforms, cell types, and infection conditions. We systematically identified 64 FDA-approved drugs as promising TB host-directed therapeutics. These high-confidence drug candidates include known HDTs such as statins (rosuvastatin, fluvastatin, lovastatin) and tamoxifen, recently validated in experimental TB models. Our prioritized candidate drugs reveal enrichment for therapeutically TB-relevant mechanisms, e.g., cholesterol metabolism inhibition and immune modulation pathways. Network analysis of disease-drug interactions identified 12 key bridging genes (including *IL-8*, *CXCR2*) that represent potential novel druggable targets for TB host-directed therapy. This work establishes transcriptome-based connectivity mapping as a viable approach for systematic HDT discovery in bacterial infections and provides a robust computational framework applicable to other infectious diseases. Our findings offer immediate opportunities for experimental validation of prioritized drug candidates and mechanistic investigation of identified druggable targets in TB pathogenesis.

## 1. Introduction

Tuberculosis (TB) is the number one infectious disease globally caused by the bacterial pathogen *Mycobacterium tuberculosis* (Mtb)^1^. Despite the availability of antibiotics, treatment of Mtb infection is prolonged and challenging due to complex features of the pathogen, including its lipid-rich, mycolic acid-containing cell wall, which restricts drug penetration and contributes to antibiotic resistance^2^. To overcome antibiotic resistance, using host-directed therapy (HDT) has emerged as a promising treatment alternative that enhances protective host immune responses^3^ rather than killing the pathogen directly. HDTs act as adjuvant therapy along with antibiotics and often multiple HDTs are combined to boost efficacy and minimize drug resistance^4–8^. However, the cost and time associated with developing entirely new HDT compounds are bottlenecks. Drug repurposing circumvents these bottlenecks by instead identifying HDT candidates from FDA-approved drugs. Computational drug repurposing accelerates drug candidate prioritization by analyzing large datasets to identify new uses for existing drugs, without requiring extensive preliminary laboratory screening^9,10^.

Transcriptome-based reversal methods have emerged as one of the primary means of computational drug repurposing; collectively referred to as ‘connectivity mapping,’ they use vast, easily accessible genome-wide gene expression responses from disease states and drug treatments^11–13^. These methods are based on the disease-induced gene expression signature—the pattern of genes turned on or off in response to disease. Connectivity mapping matches the disease signature with drug-induced signatures to find compounds that counteract the disease state and revert it to a healthy state^13^. Early efforts like the Connectivity Map (CMAP or CMAP 1.0)^11^ and the Library of Integrated Network-based Cellular Signatures (LINCS or CMAP 2.0 to emphasize the expansion upon CMAP 1.0)^12^ successfully applied this concept to prioritize therapeutics for several diseases including cancer, neurodegenerative diseases^14–19^.

Prior efforts have sought to apply transcriptomic reversal in TB, where new HDTs are urgently needed. TB-focused studies have applied connectivity mapping in relatively narrow settings, for example, using TB signatures derived from a specific profiling technology/platform and a single connectivity scoring scheme to prioritize candidate host-directed drugs. These include microarray-only meta-analysis with CMAP repurposing for TB host responses^20^, CMAP-based drug evaluation for Mtb/HIV co-infection^21^, and macrophage-focused TB signatures queried against CMAP^22^. However, analyzing a small subset of transcriptomic TB datasets with a single connectivity metric might not be representative of both biological and technical heterogeneity. Methodological differences underlying different connectivity methods often result in substantially distinct drug rankings even with the same datasets^23^. Although a benchmarking study by Gomez et al. (2024) evaluated multiple scoring strategies with various simulated data, their findings indicate that the best scoring method depends on the characteristics of the specific disease signature^24^. TB expression signatures are highly heterogeneous across laboratories, tissues, cell types, infection durations, and profiling platforms. Therefore, comparing such heterogeneous disease signatures individually to drug signatures leads to unstable predictions, because each signature often reflects a context-specific response rather than a robust, population-wise TB signal.

To navigate these challenges, we developed an integrative host-directed drug repurposing workflow incorporating six connectivity-based approaches categorized into enrichment-based^11,12^ and correlation-based metrics^25,26^. Enrichment-based methods evaluate how disease-induced expression signatures are reversed by drug-induced expression patterns, whereas correlation-based methods assess the overall similarity between disease and drug signatures. Enrichment-based approaches capture regulatory directionality of each gene in the signatures but can be sensitive to thresholding choices in defining up- and down-regulated genes^27,28^. Correlation methods can also be subject to different threshold definitions, but are simpler as they capture reversal only at the level of overall monotonic expression trends (i.e., if the disease/drug genes’ differential expression values are ranked in a similar order)^13^. Our workflow integrates both classes of methods, capturing complementary aspects of drug-disease relationships and in turn increasing the robustness and sensitivity of HDT candidate identification.

Here, we apply this new computational framework to TB. We performed a novel meta-analysis by integrating diverse transcriptomic datasets based on gene-level similarity spanning different platforms (microarray and RNAseq), infection time points, and tissue or cell types to construct a biologically representative consensus TB disease signature. This is conceptually similar to the *transfer signature* framework di Iulio et al. (2021)^29^ used to show that aggregated gene signatures derived across datasets can preserve predictive power in independent studies, e.g., across pathogens, or diseases. Building on this insight with our additional ‘weighted’ feature amplifying consistent disease signals present in the majority of diverse datasets, we hypothesized that aggregating transcriptional signatures across heterogeneous studies can yield drug reversal predictions that are more stable and biologically coherent, compared to predictions derived from individual signatures alone.

Our novel approach successfully prioritized candidates already shown to have HDT efficacy like statins ^30–33^ and tamoxifen^34^, whose mechanisms of action help tackle TB infection in macrophages. Because HDTs act by reshaping host immune responses rather than directly targeting the pathogen, their clinical utility will ultimately depend on achieving an appropriate balance between enhancing antibacterial control and avoiding excessive immune suppression, particularly in the context of combination therapy and host immune status. Despite the heterogeneity in disease-drug comparisons and data across biological and technical conditions, our TB HDT predictions show that our integrative method is a promising avenue to identify new drug target candidates. We also examined potential synergy and antagonism among current TB antibiotics and our HDT candidates, acknowledging that complementary mechanisms of action (MOAs) can enhance therapeutic efficacy^4–8^. Finally, we incorporated network-based analysis to investigate underlying disease- and drug-associated gene-gene interactions as a way to uncover mechanistic links between disease genes and known drug-interacting genes, identify shared biological pathways or processes, and prioritize candidate druggable genes based on their topological relevance within the protein-protein interaction (PPI) network. These network analyses reveal insights into shared disease-drug pathways for further investigation.

## 2. Materials and Methods

### 2.1 Disease datasets

We used all publicly available TB gene expression datasets from Gene Expression Omnibus^35^ (accessed in June 2025) with at least three TB-infected and three control samples. This collection included 10 microarray and 23 RNAseq studies (**Table S1**). We obtained uniformly processed microarray expression matrices from *refine.bio*^36^ and RNAseq expression matrices (counts) from *ARCHS4*^37^. These human gene expression datasets were heterogeneous in biological and experimental conditions, spanning a variety of five time points (in three datasets), three cell lines, 25 primary tissues, and two experimental platforms (**Figure 1a**).

**Figure 1.**
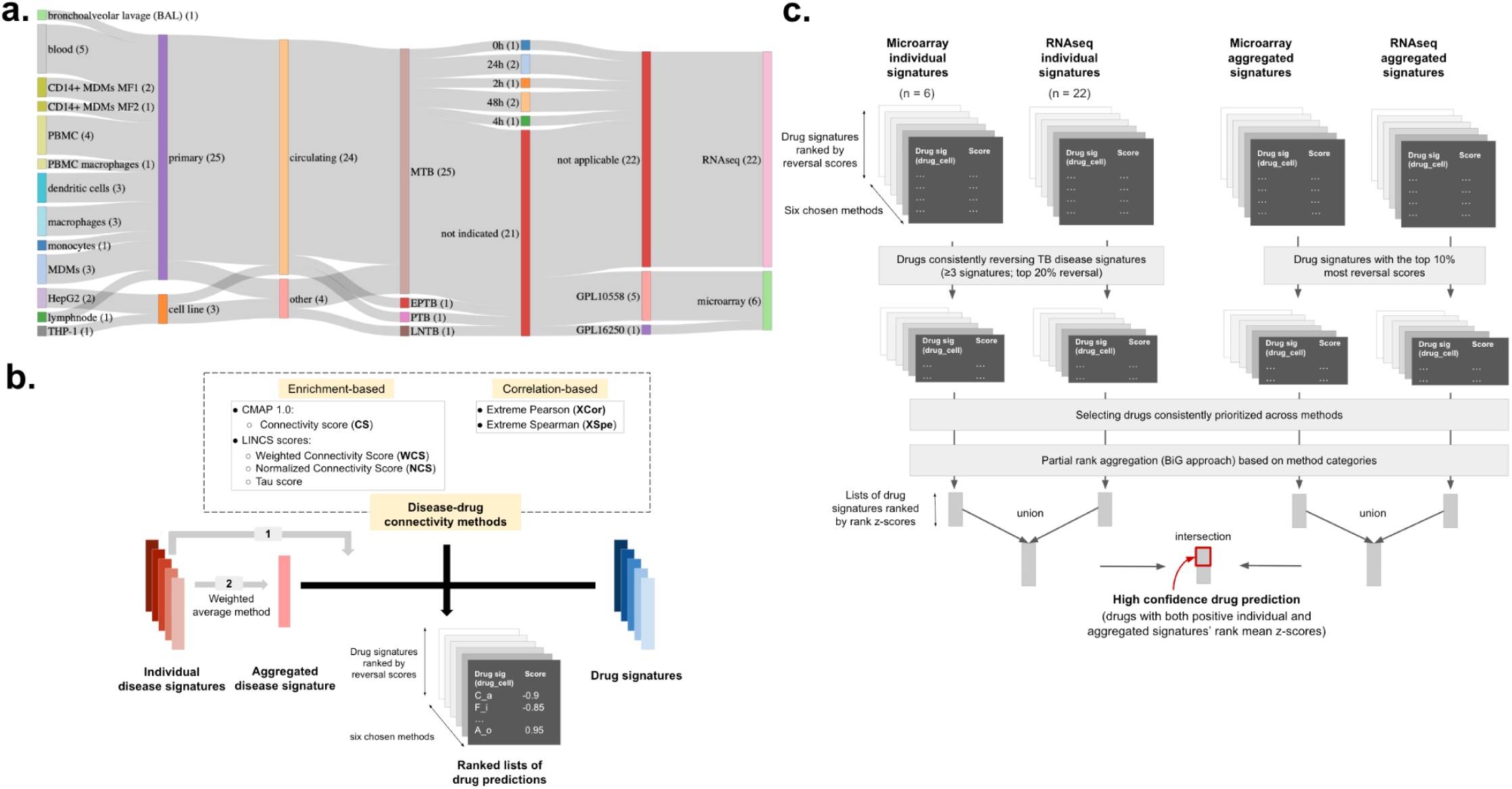
Overview of disease signature datasets and the integrative drug repurposing workflow. **(a)** Sankey diagram summarizing the biological and technical heterogeneity across the TB transcriptomic signatures. These signatures were constructed from multiple TB datasets spanning diverse biological sources, including different tissue and cell types, sample origins (blood vs. other), sample types (primary vs. cell line), TB types (PTB: pulmonary TB, EPTB: extrapulmonary tuberculosis, LNTB: lymph node tuberculosis, MTB: active TB cases without explicit “pulmonary” specification), and profiling technologies (microarray and RNAseq). **(b)** Overview of our integrative drug repurposing workflow implementing (i) individual and (ii) aggregated signatures compared against drug signatures using six connectivity scoring methods: CMAP 1.0 and CMAP 2.0 scores (WCS, NCS, τ) for enrichment-based, and Extreme Pearson (XCor) and Spearman (XSpe) for correlation-based approaches. **(c)** Drug prioritization pipeline applied to microarray and RNAseq signatures, including both individual and aggregated versions. Drugs ranked in the top 20% most negative minimum reversal scores that reversed at least three individual disease signatures were selected. From the aggregated signature predictions, the top 10% most negative-scoring drugs were selected. Subsequently, only those supported by ≥2 of 3 metric subgroups (CMAP 1.0, CMAP 2.0, correlation-based) were retained. Final drug rankings were generated using Bayesian Latent Variable Approach for partial rank aggregation; “BiG” method based on metric categories (ES- and correlation-based), and high-confidence predictions were obtained by intersecting results across signature types.

### 2.2 Constructing individual disease signatures

We performed differential gene expression analysis on microarray and RNAseq datasets using the *limma*^38^ and DESeq2^39^, respectively, comparing “infected” versus “healthy control” samples within each dataset to generate up- and down-regulated disease gene signatures (adjusted *p*-value < 0.05) and their log_2_ fold change (log_2_FC) values. The full list of disease signatures is provided in **Table S1**. We obtained six (up-/down regulated) TB-microarray and 22 TB-RNAseq signatures. These counts reflect the fact that each study can contain multiple biological conditions, and not every condition yields a gene signature.

### 2.3 Disease signature aggregation

We aggregated individual signatures (**Methods 2.2**) using *the weighted average method* outlined in **Figure S1** to generate consensus TB signatures. This aggregation step is based on the premise that differentially expressed genes consistently shared across multiple signatures are more likely to reflect core host response perturbations; therefore, we assume that signatures with higher similarity to others are likely to capture more representative disease signals. Each individual signature was assigned a weight based on its average *Jaccard similarity* to the other TB signatures. Higher similarity scores were taken to imply greater ‘confidence’ and resulted in higher weights, whereas less similar signatures were down-weighted accordingly.

For a set of *n* up-/down-regulated individual disease signatures derived from a given technology (either microarray or RNAseq data), we computed a consensus gene signature using the following steps.

1. We populated a gene membership *M* (*m* genes x *n* signatures) where *M(i,j)* is set to 0 if gene *i* is not differentially expressed in signature *j* and to gene *i*’s log_2_FC in that signature otherwise.
2. We set the overall similarity *V_j_* of each individual signature *j* equal to its average Jaccard overlap with all other signatures.
3. For each gene, summing its membership across signatures (each row in *M*) weighted by their overall similarities (*V*) resulted in its aggregated differential expression score (**Figure S2**). The final consensus signature (i.e., aggregated signature) only includes landmark genes (to match the gene coverage of the LINCS drug response signatures; **Methods 2.2**) with absolute aggregated scores greater than the 90^th^ percentile of the aggregated scores.

### 2.4 LINCS drug datasets and signatures

Currently, the LINCS database contains 1.3 million gene expression signatures of drug and small molecule perturbations tested on 3-77 different human cell lines (with the median count of six cell lines per drug), including six cancer cell lines, all generated using the L1000 assay, which measures the expression of 978 “L1000” genes^12^. We used GSE92742 LINCS level 5 data, which provided drug-perturbation signatures measured on the 978 L1000 genes (moderated z-scores aggregated at the gene-level signatures across replicates)^12^. Each drug signature is associated with specific experimental conditions: the cell line used for drug treatment, drug concentration in μM, and time point in hours. To ensure consistency with primary high-throughput compound screens in cell lines^40^, we restricted our analysis to signatures measured at 10 µM and 24 h. This condition is widely used in LINCS L1000-based connectivity analyses to enhance comparability across perturbations and cell types and represents one of the most commonly profiled settings in the compendium^41,42^. These choices limit the dataset to 8,104 compounds, including 728 FDA-approved drugs (reported by Drug Repurposing Hub^43^) across a variable number of cell lines. Limiting to FDA-approved drugs yielded 6,093 total drug expression signatures across 14 cell lines (A375, A549, ASC, FIBRNPC, HA1E, HCC515, HT29, MCF7, NEU, NPC, PC3, PHH, SKB, VCAP).

### 2.5 Finding candidate drugs using multiple scoring methods

The core principle of transcriptome-based drug repurposing is that an efficacious drug candidate should reverse disease-associated gene expression patterns (i.e., exhibit negative connectivity). Many connectivity methodologies^11,12,25,26,44,45^ quantify disease-drug negative connectivity relationships and they can be broadly categorized into enrichment score-based (ES-based) and correlation-based (pairwise similarity) methods^13^.

#### 2.5.1 Enrichment-score-based methods

ES-based methods use Gene Set Enrichment Analysis (GSEA) to calculate a disease-drug connectivity score, which captures both the distribution and membership of up- and down-regulated disease genes in a drug’s signature^11,13,46^. A more negative score indicates stronger reversal of the disease signature by the drug.

We included four ES-based methods in our analysis.

1. **CMAP 1.0 Connectivity Score (CS)** calculates a disease-drug enrichment score using a signed one-sample Kolmogorov-Smirnov (KS) test, which compares the empirical distribution of disease gene positions in the ranked drug gene list to a reference uniform distribution^11,13^.
2. **CMAP 2.0 (LINCS)** scores extend this approach using a weighted, signed two-sample KS statistic, directly adapted from GSEA, to evaluate how strongly disease genes are enriched at the extremes of a drug signature^12,13^. This scoring system includes three scores:
3. **The Weighted Connectivity Score (WCS)** provides a nonparametric measure of disease-drug similarity.
4. **The Normalized Connectivity Score (NCS)** rescales the WCS to account for variability across cell lines and drug types, enabling direct comparisons.
5. **The Tau score (τ)** further adjusts the NCS by contextualizing it within a reference distribution of connectivity scores, thereby correcting for nonspecific associations and highlighting more selective disease-drug connections.

#### 2.5.2 Pairwise-similarity-based methods

Unlike enrichment-score-based methods, pairwise-similarity correlation-based approaches capture the degree and direction of overall similarity between the expression patterns of disease and drug; a negative correlation suggests a potential reversal effect. Here, we implemented two such methods, denoted as **XCor** and **XSpe** ^13,25,26^. These two metrics are based on the correlation between two ranked sets of values; in our case, correlations are between differential gene expression values (log_2_FC) for a pair of disease and drug signatures. XCor and XSpe calculate the Pearson and Spearman correlation^47^, respectively, between log_2_FC values of significantly differentially expressed genes (adjusted *p*-value < 0.05) shared between the disease and drug signatures.

### 2.6 Prioritizing top drug candidates

For each drug, we calculated the minimum reversal score across multiple cell lines to reflect the best reversal scenario of the drug across all available cell lines and used this cell line-agnostic score in downstream drug prioritization.

#### 2.6.1 Selection of drugs reversing the majority of individual disease signatures

To identify strong drug candidates, for each of the six connectivity methods, we prioritized the top 20% of the drugs with the strongest negative connectivity scores, then selected drugs reversing at least three individual disease signatures at the 80^th^ percentile reversal threshold.

#### 2.6.2 Selection of drugs reversing aggregated signatures

To prioritize drugs based on the aggregated disease signatures, per connectivity method, we selected the top 10% of drugs with the most negative connectivity scores.

#### 2.6.3 Method-wise filtering and rank aggregation

All analyses described above, from disease signature construction to drug prioritization using each of the six connectivity metrics, were performed independently for microarray and RNAseq datasets, and separately for individual and aggregated disease signatures. This resulted in four separate drug prioritization pipelines: (i) individual microarray signatures, (ii) aggregated microarray signatures, (iii) individual RNAseq signatures, and (iv) aggregated RNAseq signatures. We next sought to consolidate these results by performing method-wise filtering and rank aggregation within each pipeline to identify robust top candidates.

To ensure robust drug prediction results across different connectivity measures, the drugs appearing only in one of the three method categories (CMAP 1.0, CMAP 2.0, and correlation-based) were excluded. Consequently, analyzing results from microarray and RNAseq separately, the remaining drugs in the lists from both individual and aggregated disease signatures were then summarized into two drug lists (one per signature type) using the rank aggregation method ‘BiG’^48,49^, which has been shown to provide robust aggregation performance regardless of heterogeneity in method effectiveness across the input lists^48^. Here, we aggregated drug lists by considering the ES-based and correlation-based connectivity scores (**Figure 1c**) as two sources of heterogeneity (referred as “platform bias” in Li et al., (2019)^49^). This step resulted in four rank-aggregated drug lists: microarray or RNAseq and individual or aggregated.

#### 2.6.4 Finalization of high-confidence predicted drugs

To identify a robust set of candidate drugs, we first took the union of microarray and RNAseq rank-aggregated results (each derived across multiple connectivity metrics) separately for the individual and aggregated TB signatures and then computed their intersection (**Figure 1c**). The high-confidence drugs on the final list were ranked by their combined mean-rank z-scores (*z*_*combined*_ ; **Equation 1**). We further prioritized a subset of 64 top candidates by selecting those with positive *z*_*indiv*_ and *z_aggr_* (**Equation 1**), indicating consistent and above-average reversal across both individual and aggregated TB signatures (**Table S2**).

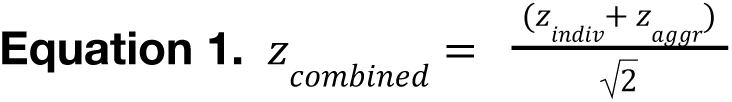

where *z* _*combined*_ is combined mean-rank z-score,

*z _*indiv*_* and *z* _*aggr*_ are the mean-rank z-score of drug predictions derived from individual and aggregated signatures, respectively.

Both *z* _*indiv*_ and *z* _*aggr*_ are calculated as the z-score of the average rank of each drug in the microarray and RNAseq rank-aggregation result lists.

### 2.7 Disease pathway enrichment analysis

#### 2.7.1 Biological process enrichment calculation

To delineate perturbed biological processes underlying TB, we performed pathway enrichment analysis on the individual disease signatures from both microarray and RNAseq (from 2.2 and 2.3) to identify commonly enriched Gene Ontology Biological Process (GO:BP) terms^50^. We restricted to GO:BP terms with sizes of at least five genes and most 200 genes to ensure ample statistical power and reduce statistical bias towards general terms. The enriched GO:BP terms were identified using the *ClusterProfiler* R package^51^, using genes annotated to GO terms present in any of our disease datasets from either of the technologies as the background genes. We quantified the level of enrichment as the inverse hyperbolic sine (*asinh)* of the ratio between the observed overlap of input genes associated with a given GO term and the expected overlap by chance (**Equation 2**). The pathways with high enrichment serve as a proxy for robust, cross-study biological signals.

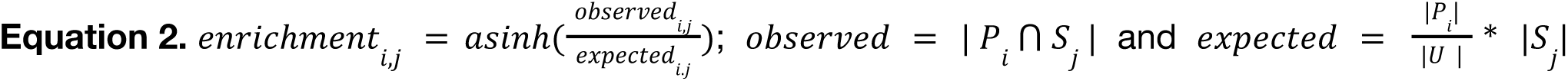

where *P_i_* is the gene set that is part of pathway *i*,

*S_j_* is the differentially expressed gene set of signature *j*,

and *U* is the universe gene set; measured genes across all signatures.

#### 2.7.2 Evaluation of the effects of confounding variables in disease pathway enrichment via statistical tests and highly variable enriched pathways/processes

We first examined consistency of pathway enrichment across our individual TB signatures by performing variance analysis on all pathways’ enrichment scores (**Equation 2**) of the individual signatures. Variance was computed to assess whether the pathways are conserved in most signatures (low variance) or vary across signatures (high variance) (**Figure S3**).

Next, we investigated whether observed differences in pathway enrichment could be explained by specific study-level metadata — (i) baseline type (primary sample vs. cell line), (ii) tissue origin (circulating vs. other), and (iii) profiling technology (microarray vs. RNAseq) — rather than true disease signals. We followed the pathway analysis results (**Methods 2.7.1**) with visualization of the enrichment distributions of the most variable GO:BP terms across the individual signatures stratified by metadata (**Figure 2a,b**). We derived clusters based on semantic similarity, using the *GOSim* R package^52^, from GO:BP terms whose enrichment variances were above the 90^th^ percentile across individual signatures. We selected the highest-variance GO:BP term in each cluster as the cluster representative for visualization in **Figure 2** (panels **a** and **b**).

**Figure 2.**
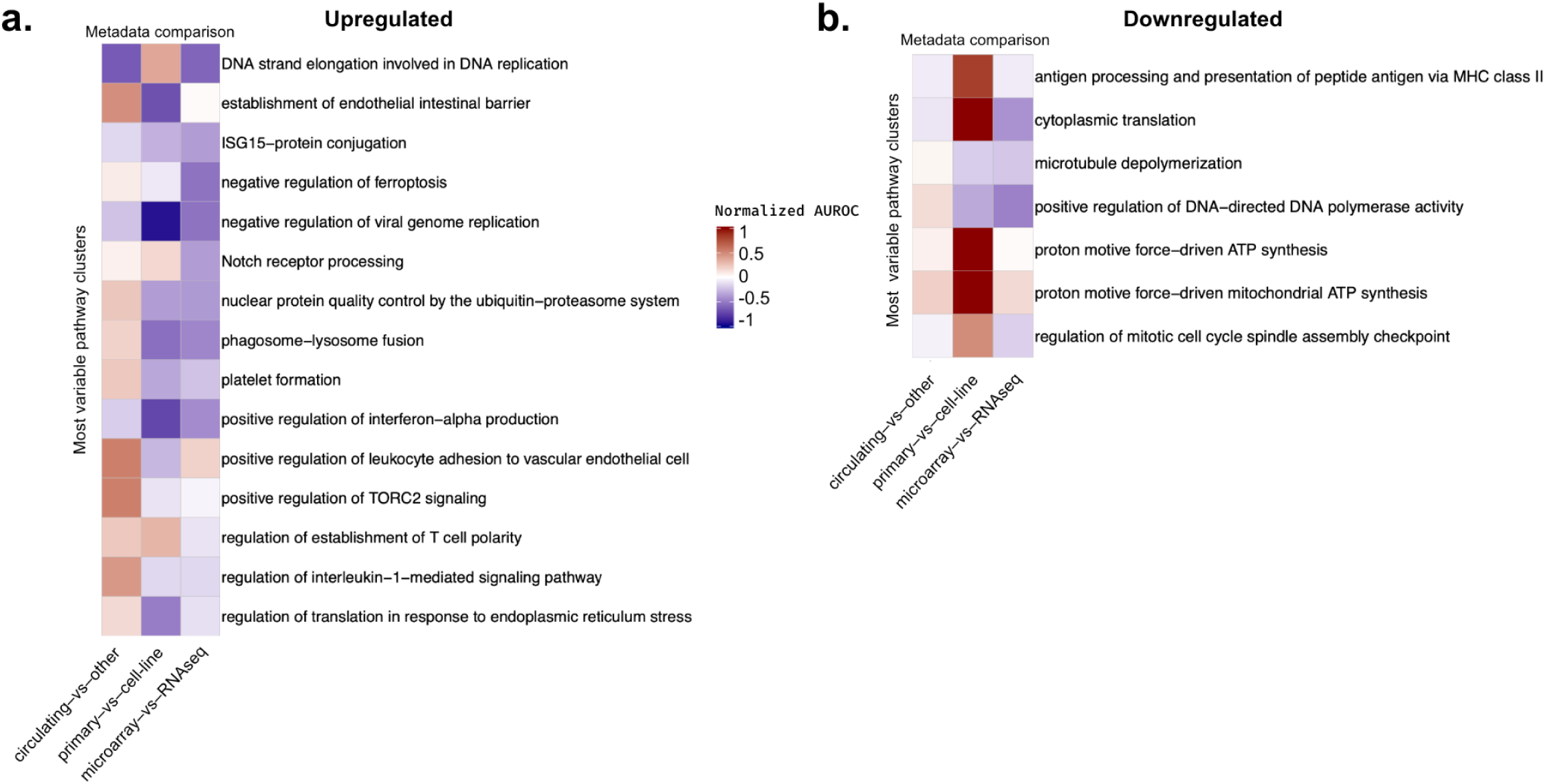
Heterogeneity of TB responses through the lens of highly variable biological pathways and processes. Normalized AUROC scores across the top highly variable cluster representative **(a)** up- and **(b)** down-regulated GO:BP by metadata data contrasts. The red-to-purple scale indicates normalized enrichment (i.e., how these highly variable GO:BP terms are enriched in group one (circulating, microarray, or primary; red) or group two (other, RNAseq, or cell line; purple)) from highest to lowest. The white cells represent no enrichment of the GO:BP terms.

For each metadata category with at least three signatures, we applied a Mann-Whitney U test to assess whether the enrichment distribution across all individual signatures of any pathway differ significantly between metadata-specific groups (e.g., circulating vs. non-circulating tissue). We then quantified the heterogeneity using normalized area under the receiving operator characteristic (AUROC) curve, a rank-based measure derived from the Mann-Whitney U statistic used in our statistical testing ranging from -1 to 1 (**Equation 3**). Positive *norm*(*AUROC*) values indicate enrichment of a GO:BP term in the first group (e.g., circulating), while negative values reflect enrichment in the second group (e.g., other). We further visualized normalized AUROC distributions for highly variable GO:BP terms stratified by metadata groups to highlight terms specifically more enriched in one metagroup than the other (**Figure 2c**).

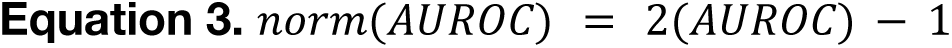

*U*_1_ is Mann-Whitney U statistic of the metadata group 1

(restricted to only group 1 to maintain directionality),

*n*_1_ is the number of signatures belonging to the metadata group 1, and

*n*_2_ is the number of signatures belonging to the metadata group 2.

#### 2.7.3 Evaluation of pathway-level agreement between individual and aggregated signatures via deviation analysis

To evaluate how well the aggregated disease signatures reflect the biological signals captured in individual signatures, we classified enriched GO:BP terms as “dominant” (enriched in >90% of the individual signatures) and “non-dominant” (otherwise). Next, we analyzed which TB signals are well-captured by aggregated signatures or deviate from host responses in individual signatures. We summarized each pathway’s enrichment strength by computing the median enrichment across all individual signatures where that pathway appeared. We then quantified deviation by computing squared differences between the aggregated gene ratio and this median, normalized by the number of individual signatures (**Equation 4**). The intuition is that a pathway that is well-captured by aggregated signatures would correspond to a deviation score close to zero (**Figure 3**).

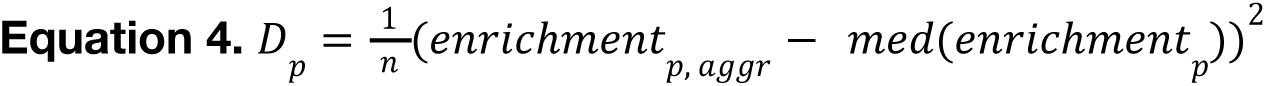

**Figure 3.**
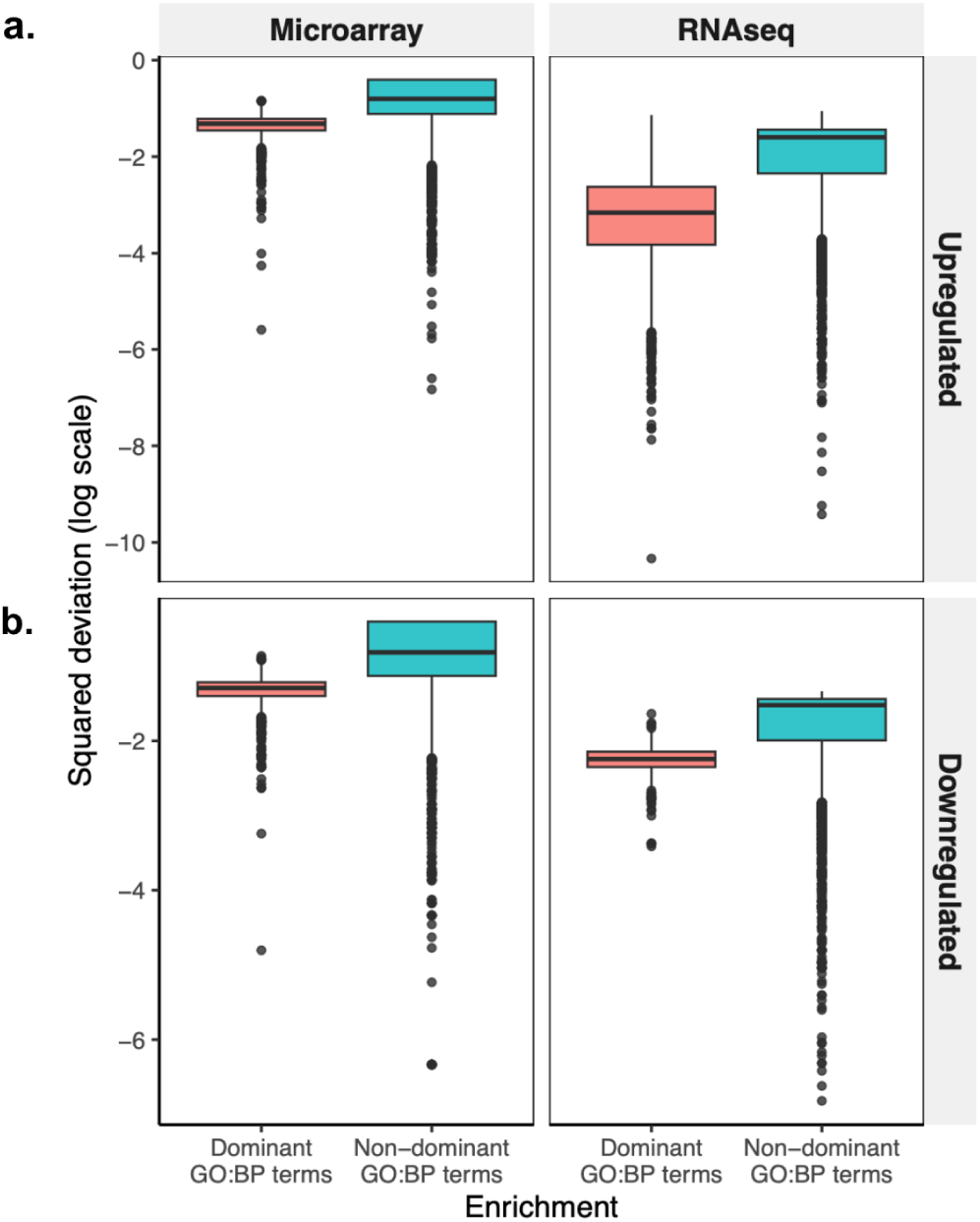
Evaluation of pathway-level deviation between aggregated and individual signatures across technologies. **(a)** Upregulated and **(b)** downregulated GO:BP pathways are grouped by their prevalence across individual TB signatures (<90% vs ≥90% enriched) and stratified by transcriptomic technology (microarray and RNAseq). For each pathway, squared deviation quantifies the difference between enrichment in the aggregated signature and the median enrichment across individual signatures. Boxes indicate the interquartile range with median; whiskers denote 1.5 × IQR. The y-axis is shown on a log scale. Statistical significance between groups was assessed using a Mann–Whitney U test (FDR-adjusted *p-value* < 0.05).

where *D* _*p*_ is deviation score for pathway *p*,

*n* is number of individual signatures,

*enrichment* _*p*,*aggr*_ is enrichment of pathway *p* in aggregated signature, and

*med*(*r_p_*) is median enrichment of pathway *p* across all *n* individual signatures.

### 2.8 Constructing cholesterol- and cytokine-related disease-drug pathway networks

To investigate shared biological mechanisms between predicted HDTs and TB-related disease genes, we constructed two pathway-relevant networks focused on cholesterol metabolism and cytokine signaling in human macrophages. These pathways were selected based on (i) the most frequent occurrence of HMGCR inhibitors (statins), a well-known drug class, among the top-ranked predicted drugs and (ii) the central role of cytokine signaling in macrophage immune responses against Mtb^53^.

For each drug in the final prediction list, we included its drug-interacting genes with interaction scores in the top 90^th^ percentile obtained from the Drug-Gene Interaction Database (DGIdb)^54^ , and the top three most positively and negatively perturbed genes based on GSE92742 LINCS level 5 expression data (24-hour, 10 μM conditions)^12^. Drugs were linked based on shared enriched pathways identified via Drug Set Enrichment Analysis (DSEA) implemented in the *signatureSearch* package^40^. Then, we restricted the network to the subset of drugs annotated to either cholesterol- or cytokine-associated pathways. To expand each network, we integrated high-confidence inferred protein-protein interactions (confidence score ≥ 0.7) from the STRING database^55^. Finally, we used the intersection of each network with aggregated TB disease gene signatures to identify shared components between disease-related genes and drug-associated molecular effects.

### 2.9 Building disease-drug gene/pathway networks

To identify potential mechanistic links between TB disease genes and predicted drug targets, we constructed disease-gene/pathway-drug networks using the STRING network^55^. The disease-associated genes were derived from our aggregated TB signature, combining upregulated and downregulated genes that consistently appear in both microarray and RNAseq datasets. This resulted in a total of 16 disease genes, defined as *source nodes*. For each predicted drug, we obtained known drug-interacting genes (i.e., *target* nodes) from DGIdb^54^, selecting only those genes with interaction scores in the top 90^th^ percentile. Additionally, the top four drug-perturbed genes (top two up-/down-regulated genes) were extracted from LINCS level 5 drug signature data for every drug and incorporated as complementary target nodes. For each disease gene/drug-interacting gene pair, the shortest paths within the STRING network were calculated using Dijkstra’s algorithm and A*^56^. Then we computed the average betweenness centrality (BC) score per node for all nodes along the pre-computed shortest paths. A gene’s BC quantifies a gene’s importance in facilitating information flow in the PPI network, prioritizing genes that are likely to serve as key intermediates between disease pathways and drug mechanisms of action. We selected the top 2% of paths with the highest average BC scores per source-target gene pair. To ensure that the predicted novel druggable genes function within specific TB-associated pathways, rather than generically, we explicitly marked genes that appear in the top 20% of drugs with highest BC scores as hub genes in the network visualization (**Figure 4**).

**Figure 4.**
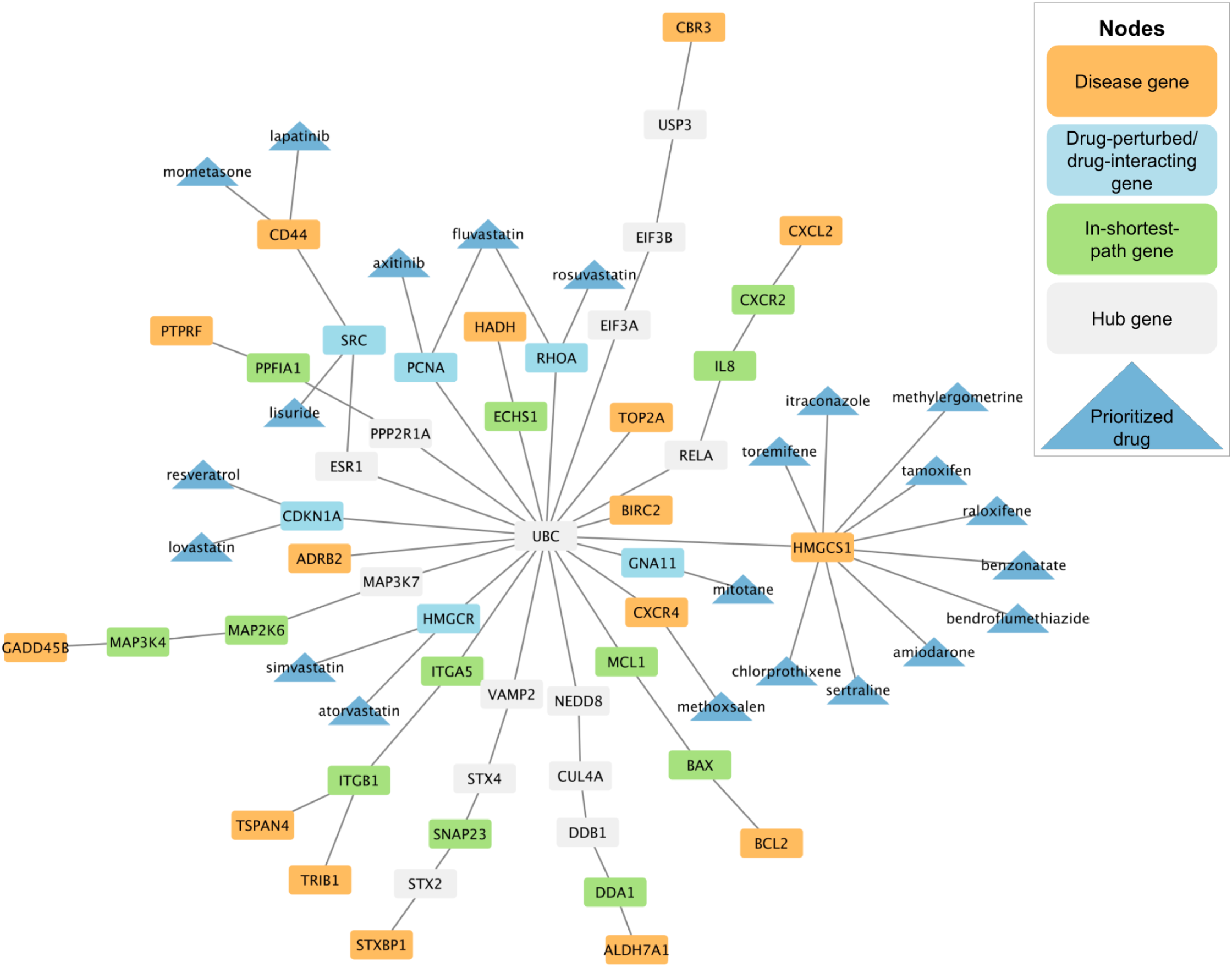
Disease-drug-target network identifies key linking genes and prioritized drug candidates. Network visualization showing shortest-path connections between TB host genes perturbed by active TB infection (orange), known drug-interacting or drug-perturbed genes (light blue), and prioritized drugs (dark blue) through intermediate in-path genes (green) within the STRING genome-wide host protein-protein interaction network. Hub genes (grey) are the top 20% genes with highest betweenness centrality. This analysis highlights 12 key in-path genes including *IL-8, CXCR2, MCL-1, BAX, SNAP23, ITGB1, ITGA5* that serve as mechanistic bridges between TB-perturbed genes and known drug targets. Prioritized candidates (dark blue nodes) include known TB-relevant HDTs (e.g., rosuvastatin, tamoxifen) and novel predictions (mitotane, amiodarone, sertraline). These network-level insights support the biological relevance of drug predictions and suggest new avenues for host-directed TB therapeutics.

### 2.10 Predicting potential drug mechanism of action pairs showing synergistic and antagonistic effects

We sought to identify synergistic drug combinations for enhanced efficacy based on key MOAs associated with our predicted TB HDT candidates. We first took (i) our prioritized HDT drug candidates from both microarray and RNAseq TB datasets and (ii) current TB antibiotics^57^ and their corresponding MOAs. For each drug pair among the predicted candidates, we retrieved corresponding synergy scores derived from high throughput screening reported in *DrugCombDB* (retrieved on 07/24/2024)^58^.

We defined synergistic drug pairs as those reported “synergy” by ≥3/4 methods in *DrugCombDB* (ZIP^59^ , Bliss^60^ , Loewe^61^ , and HSA^62^) and antagonistic pairs as those reported “antagonism” by ≥3/4 methods. Then at the MOA-level, we computed the fractional synergistic score to measure how likely each MOA pair demonstrates synergy or not based on the drug-level information where we ensured that each drug pair only contributes at most once to a given MOA pair.

### 2.11 Comparison of “healthy” and “untreated” baselines

To assess if the tissue/cell type used for profiling the disease and the drug should be accounted for when making reversal predictions, we evaluated the similarity between healthy control disease samples and untreated drug cell line profiles for biologically meaningful correspondence.

We selected L1000 Level 3 profiles measured at 24-hour, applied quantile normalization, and aggregated them across each cell line using either the mean or Stouffer-like weighted mean (pairwise correlations among replicates to derive weights that emphasize internally consistent samples) to create the drug baseline profiles. The control disease samples were obtained from the microarray and RNAseq TB datasets, filtered to match the landmark genes, and quantile transformed to match the distribution of the drug baseline data. To compare these disease and drug baselines pairwise, we applied four correlation metrics—Pearson, Spearman, Rank-biased overlap (RBO)^63^, and Lasso regression coefficients—that capture different aspects of concordance of gene expression values. Pearson and Spearman measure linear and monotonic associations, respectively, RBO emphasizes top-ranked gene agreement, and Lasso regression incorporates sparsity to retain only predictive relationships.

## Results

### 3.1 Systematic prediction of host-directed therapeutics from TB transcriptomes

Our integrative computational framework reconciles the multiple sources of heterogeneity inherent in public transcriptomics datasets, including differences in profiling technologies (microarray and RNAseq), biological contexts (tissues, cell types, cell lines, and time points), and drug scoring methods, to enable robust drug prioritization (**Figure 1a**). We started with 10 microarray and 23 RNAseq studies, used standard differential expression analysis to obtain 6 microarray and 22 RNAseq TB disease signatures (**Methods 2.1, 2.2; Table S1**), and combined them using a weighted Jaccard similarity approach to derive an aggregated TB signature capturing shared host responses across datasets (**Methods 2.3**). For both individual and aggregated signatures, we performed disease-drug comparisons with LINCS drug signatures (**Methods 2.4**) using ES-based, and correlation-based methods (**Figure 1b**). We integrated these drug results, retaining drugs that appeared consistently across both microarray and RNAseq signatures, and derived 214 candidate HDTs (**Methods 2.5, 2.6; Figure 1c; Table S2**). We further narrowed our high-confidence drug list to 64 candidates that showed consistent and above-average reversal across both individual and aggregated TB signatures (using positive combined z-scores, **Table S2**). These candidates include both known TB-relevant drugs like statins^30,76^, tamoxifen^34^, and novel predictions such as clonidine previously used for hypertension (**Tables 1, S2**).

### 3.2 Host responses to TB vary by biological and technical factors

Next, to examine how technical factors contribute to differences in TB responses recorded in transcriptomics studies (**Table S1**), we captured gene and pathway activity across signatures. First, we observed that TB signatures cluster primarily based on both biological and technical factors (**Figure S1**).

To disentangle sources of heterogeneity, we tested whether most variable pathways showed enrichment differences when signatures were grouped by metadata (biological or technical factors): (i) tissue type: circulating vs. other, (ii) source of origin: primary sample vs. cell line, and (iii) gene expression technology: microarray vs. RNAseq. We quantified term enrichment using a normalized *AUROC* score (**Methods 2.7.2; Equation 3**) across each metadata contrast (e.g., circulating vs. other; **Figure 2**; top variable GO:BP shown in **Figure S4**). In summary, our results convey that both biological and technical variations contribute to the differences in host response patterns to TB. We highlight a few key findings below.

#### 3.2.1 Pathway enrichment of circulating and non-circulating TB signatures reflects systemic versus local TB responses

The most variable GO:BP terms, by definition, are sparsely enriched across individual signatures (**Figure S4a,b**). Circulating signatures (e.g., whole blood, PBMCs; **Figure S4a,b**) are not enriched for any most variable terms shown in **Figure 2** in the Mann-Whitney U test (**Methods 2.7.2; Table S3)**. However, their most upregulated terms mostly exhibit positive *norm*(*AUROC*) values (**Figure 2**) showing that circulating TB signatures were enriched for processes related to interferon and interleukin signaling, leukocyte adhesion, phagolysosomal maturation, and protein quality control which are involved in systemic immune activation^64,65^. DNA replication-associated processes and canonical antiviral interferon-stimulated gene (ISG) effector pathways were enriched in BAL- and cell-line TB signatures and not circulating cells, reflecting cell stress and interferon responses at the site of infection (**Figure 2**; **Table S3)**^66–68^.

For the most variable downregulated processes, differentiation of enrichment between circulating and non-circulating signatures was weak. This is reflected in the low *norm(AUROC)* values (**Figure 2**), implying these processes might be similarly represented in both circulating and other tissue compartments. Despite the weak signals, the results still convey different responses at systemic and local levels. Many terms enriched in circulating signatures, such as microtubule dynamics, spindle checkpoint regulation, DNA polymerase activity, and mitochondrial ATP synthesis, reflect broad changes in cell cycle control, cytoskeletal organization, and cellular energetics rather than compartment-specific biology. In contrast, BAL and cell line signatures captured downregulation of antigen processing and presentation via MHC class II and cytoplasmic translation expected as part of cell activity in BAL^69^ and stronger cellular transcriptional responses *in vitro* cell line systems^70^ (**Figure 2**).

#### 3.2.2 Microarray and RNAseq capture distinct pathway-level signals in TB

We compared enrichment patterns between microarray and RNAseq signatures. Sample size imbalance reduced statistical power; as expected, highly variable GO:BP terms did not visibly separate by technology (**Figure 2a,b**), and Mann-Whitney tests identified no significantly differentiating terms. However, *norm*(*AUROC*) scores revealed technology-specific enrichment patterns (**Figure 2**). Microarray signatures showed higher *norm*(*AUROC*) values for a subset of the most variable upregulated terms (**Figure 2a**); DNA strand elongation in replication, Notch receptor processing, and regulation of establishment of T cell polarity. Most immune- and stress-related terms showed stronger enrichment in RNAseq signatures.

Among downregulated terms (**Figure 2b**), only microtubule depolymerization and DNA polymerase regulation showed stronger enrichment in microarray signatures. Antigen presentation, protein synthesis, mitochondrial energy metabolism, and mitotic cell cycle pathways were more strongly enriched in RNAseq. These platform-specific differences reflect underlying technical characteristics: microarrays emphasize well-annotated, moderately-to-highly expressed genes, while RNAseq captures broader immune, metabolic, and stress programs through greater dynamic range and sensitivity to low-abundance transcripts^71,72^.

#### 3.2.3 Primary and cell-line TB signatures highlight different metabolic and translational responses

Next, we compared primary samples and cell lines. Despite inconclusive clusters (**Figure S4a,b**), *norm*(*AUROC*) scores (**Figure 2**) and Mann-Whitney statistical tests (**Table S4**) revealed pathway-level differences. Among upregulated pathways (**Figure 2a**), primary signatures were enriched for tissue interaction and immune cell trafficking, including endothelial barrier function, leukocyte adhesion to vascular endothelium, and TORC2 signaling. Cell line signatures were instead enriched for DNA replication, Notch receptor processing, interferon-related responses, protein quality control, and phagosome-lysosome fusion. Among downregulated terms (**Figure 2b**), primary signatures were enriched for mitochondrial ATP production, while cell line signatures included antigen presentation, cytoplasmic translation, cytoskeletal dynamics, DNA polymerase regulation, and mitotic checkpoint control.

Cell line-specific pathways included mitochondrial organization and metabolic stress responses (upregulated; **Table S4a**), and translational control, RNA handling, and developmental programs (downregulated; **Table S4b**). These patterns indicate that primary samples reflect tissue-contextual and metabolic programs, whereas cell-line models emphasize proliferative, stress-associated, and biosynthetic responses characteristic of *in vitro* infection^73,74^, consistent with studies comparing cell lines to their tissues of origin^75^. Overall, these findings suggest that TB transcriptomic signatures reveal heterogeneous host response, often confounded by biological and experimental factors, and profiling technology. We, therefore, need to factor this in while interpreting TB host responses, performing downstream analyses such as aggregation, and repositioning drugs.

### 3.3 Aggregated signatures capture dominant biological signals across individual signatures

Given the heterogeneity across individual TB signatures (**Result 3.2**), we tested whether aggregation could summarize dominant, biologically relevant TB signals. We combined individual signatures into one aggregated signature for up- and down-regulated gene sets independently, using a weighted average approach with *Jaccard* similarity as a proxy for ‘confidence’ (**Methods 2.3**; **Figure S1a,b**). Signatures with higher similarity to other signatures received higher weights, while dissimilar signatures were downweighted. The goal was to capture dominant TB perturbations consistent across our 28 individual signatures while downweighting signals from potentially confounding variables associated with individual signatures, such as experimental platform, tissue/cell type, and infection time point.

To evaluate whether aggregated signatures captured major biological processes, we examined aggregation coverage of dominant GO:BP pathways enriched across individual datasets (**Methods 2.7.3**). All “dominant” pathways (conserved in ≥90% of signatures) showed significantly lower deviation scores than “non-dominant” pathways (FDR-adjusted *p-value* < 0.05; **Figure 3**; **Table S5**), indicating successful recapitulation of conserved biological signals underlying TB host response.

Non-dominant pathways showed substantially higher deviation scores, indicating poor retention in aggregated signatures. These pathways include diverse metabolic, biosynthetic, and cellular maintenance processes: nucleotide and amino acid biosynthesis, mitochondrial and lipid metabolism, protein folding, transport, and oxidative stress regulation (**Figure S5**). These processes showed enrichment only in specific signature subsets depending on biological context, and were therefore downweighted during aggregation.

These findings indicate that our aggregation method captures globally conserved host responses to TB, but not context-specific signals that depend on cell or tissue type.

### 3.4 Predicted candidate HDTs with literature evidence against TB

Building on aggregated signatures’ ability to capture dominant TB signals, we developed an integrative workflow to repurpose candidate HDTs (**Methods 2.5–2.6**). We evaluated whether top predicted candidates included known or emerging HDTs with anti-TB efficacy and used published literature to assess biological relevance and provide external validation.

As proof-of-principle, our workflow successfully prioritized known HDTs for TB, including tamoxifen^34^ and statins^30,76^: atorvastatin^77,78^, fluvastatin^79^, lovastatin^80^, and rosuvastatin^81,82^. The candidate tamoxifen has shown activity against TB infection both *in vitro* and *in vivo* by enhancing lysosomal activity and increasing Mycobacterial delivery to lysosomes, reducing bacterial survival in cell and animal models^34^ (**Table 1**).

**Table 1.**
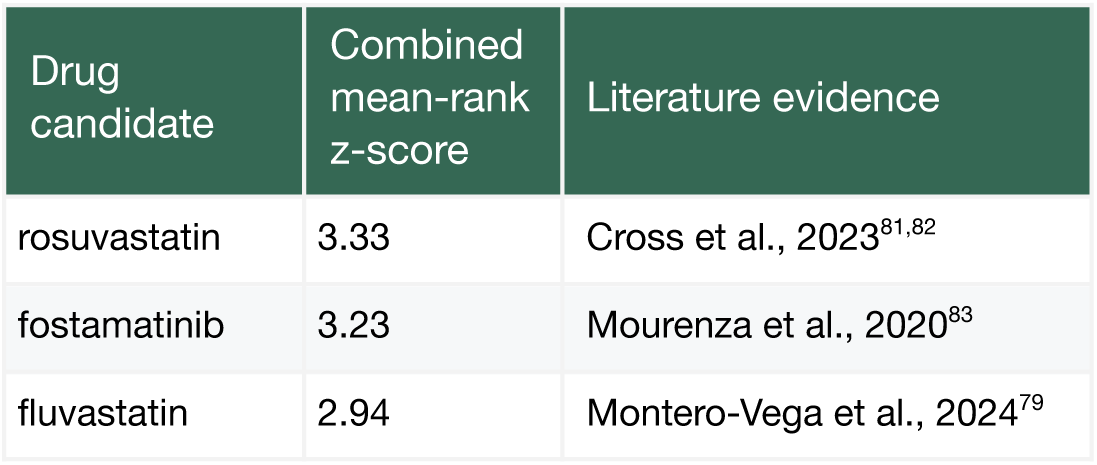
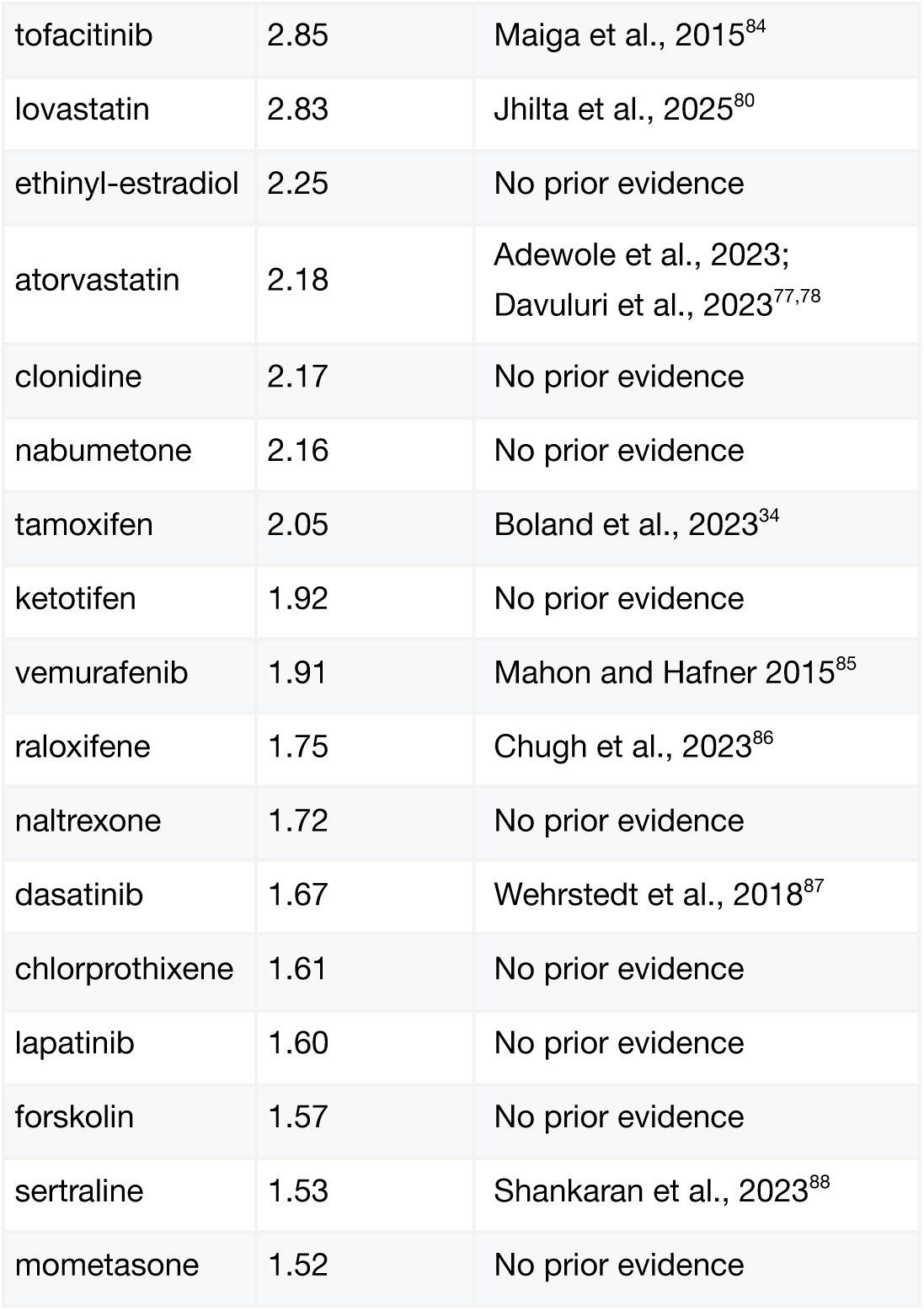
Top 20 high-confidence drug candidates. Drug candidates were prioritized using combined mean rank z-scores, which reflect the overall ranking of each drug across predictions from both individual and aggregated TB signatures using six complementary connectivity scoring methods. The table shows the top 20 drug candidates, along with literature evidence supporting their potential as HDTs against TB. “No prior evidence” indicates no published support for anti-TB activity. See **Table S2** for the full list of predicted candidates.

Top predicted candidates showed distinct MOAs, including HMGCR inhibitors (statins) and PDGFR pathway inhibitors (axitinib, dasatinib, trapidil), were frequently observed across microarray and RNAseq datasets (**Table 2; Figure S8**). We investigated molecular mechanisms underlying the 28 TB signatures (**Table S2**) to identify technical and biological signals driving our predictions.

**Table 2.**
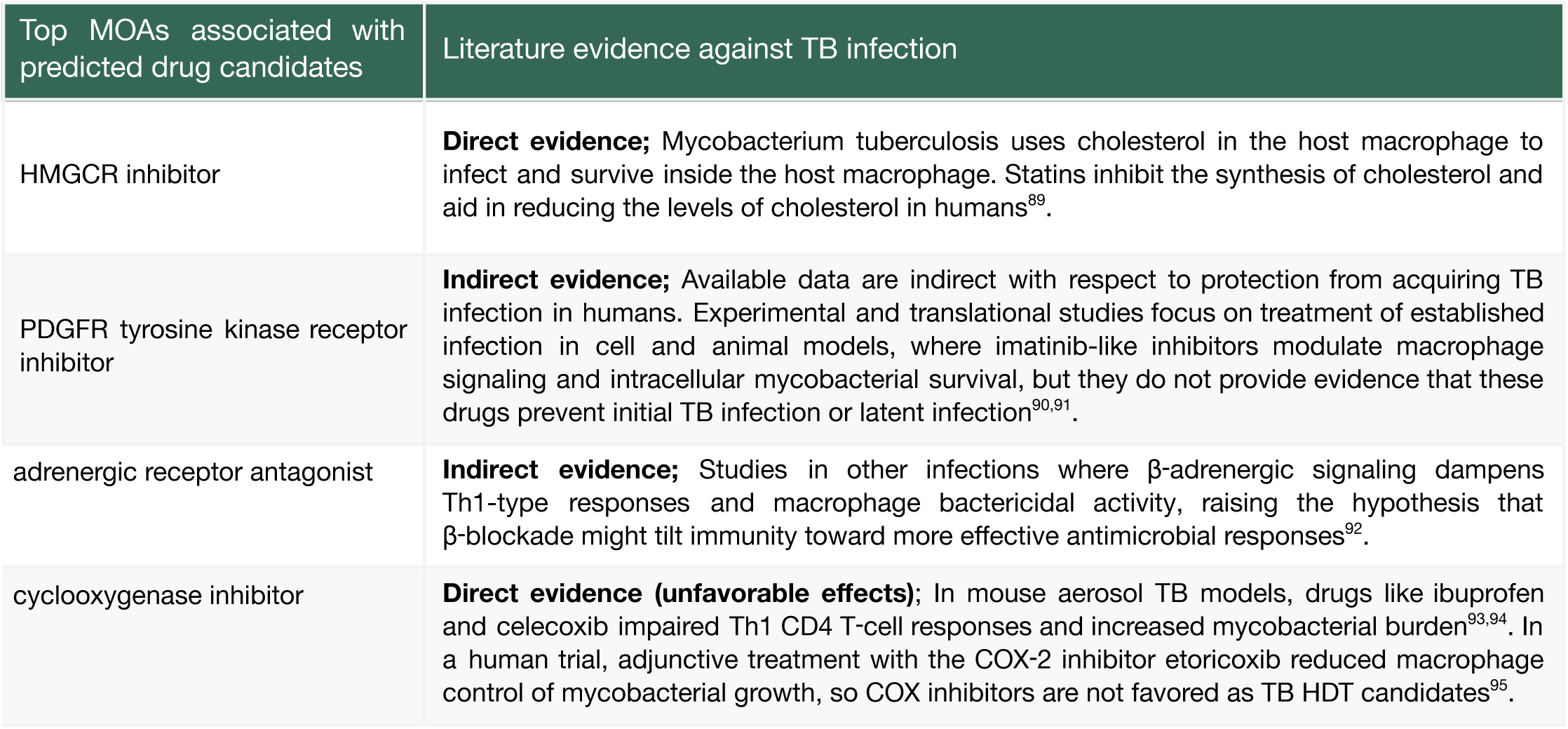
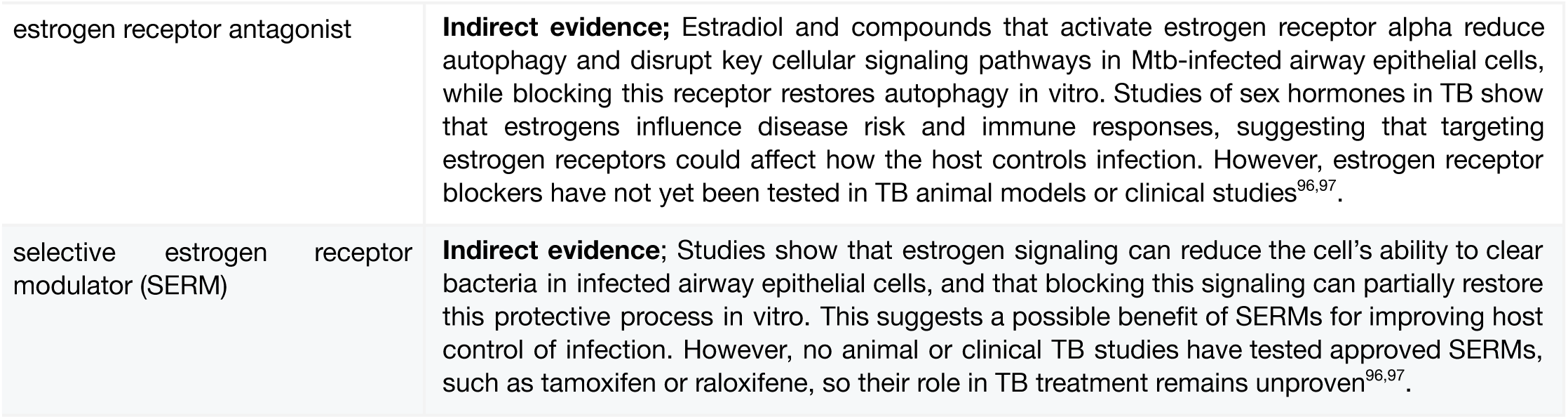
Mechanisms of action (MOAs) associated with predicted drug results. MOAs linked to top-ranked drug candidates with literature evidence describing their relevance to TB infection or host-directed mechanisms. References highlight prior studies supporting the therapeutic relevance of these MOAs against Mtb.

### 3.5 Shared disease-drug pathways involving cholesterol and cytokine signaling

To examine how our predicted drug and disease genes converge on cholesterol- and cytokine-related biological processes, we constructed two networks for cholesterol- and cytokine-related pathways (**Figure S7**). For each network, we incorporated known targets of the predicted drugs and drug-perturbed genes (the three most differentially expressed up- and down-regulated genes from LINCS drug signatures). We connected drugs based on shared pathway enrichment using DSEA (**Methods 2.8**), and expanded networks with high-confidence inferred interactions from the STRING database.

Both networks revealed a shared node, *HMGCS1*, linking disease- and drug-perturbed genes. *HMGCS1* encodes a key enzyme in the mevalonate pathway, which intersects inflammatory and metabolic signaling, indicating convergence of predicted HDTs on cytokine and cholesterol pathways (**Figure S7**).

### 3.6 Novel druggable gene targets and high-confidence TB HDT candidates via a network analysis

Our integrative transcriptome-based drug prioritization relies on global reversal patterns between disease and drug signatures. Network analysis provides additional gene-level insights by identifying *linking genes* that bridge disease-perturbed genes and known drug targets. These in-path genes reveal mechanistic connections between disease and drug actions, revealing new therapeutic targets. Using this approach in **Methods 2.9**, we identified 12 key in-path genes linking TB disease genes and drug targets: *PPFIA1, ECHS1, IL-8, CXCR2, MCL-1, BAX, DDA1, SNAP23, ITGB1, ITGA5, MAP3PK4, MAP2K6* (**Figure 4**; **Table 3**).

**Table 3.** Identified potential druggable genes with literature evidence. Key host genes identified in the disease-drug-target network as mechanistic bridges (in-path genes) connecting TB-perturbed genes and known drug targets. Each gene has literature evidence for involvement in host immune response or cellular processes relevant to TB infection, serving as potential targets for host-directed TB therapeutics.

| Identified in-path genes | Literature evidence against TB infection |
| --- | --- |
| PPFIA1 | **Indirect evidence;** PPFIA1 is linked to crucial cellular processes and immune regulatory pathways relevant to TB pathogenesis, specifically immune cell trafficking and granuloma formation<sup>102</sup>, cell survival/apoptosis<sup>103</sup>, initiating inflammation<sup>103</sup>. In mouse models with chronic myeloid leukemia, targeting PPFIA1 effectively suppressed the self-renewal and promoted the apoptosis of leukemic stem cells<sup>104</sup>. |
| ECHS1 | **Indirect evidence;** ECHS1 acts as a negative regulator of the cGAS/STING/Type I IFN signaling pathway through metabolic reprogramming<sup>105</sup>. This pathway is critical for recognizing intracellular pathogens and inducing autophagy for bacterial clearance<sup>106</sup>. Literature documents inhibition of ECHS1 as a viable strategy to boost innate immune response<sup>98</sup>. |
| IL-8 | **Direct evidence;** IL-8 is a key molecular marker of the immune response to Mtb<sup>98</sup> and responsible for recruiting T-cells<sup>99</sup>. Literature suggests a dual strategy regarding IL-8, indicating that the optimal approach depends heavily on the stage of the infection due to IL-8's beneficial early functions and potentially detrimental late-stage effects<sup>100</sup>. |
| CXCR2 | **Direct evidence;** CXCR2 is identified as a key regulator of neutrophil subpopulations and is crucial for TB immunopathogenesis<sup>101</sup>. Treatment with a specific CXCR2 inhibitor in TB-susceptible mouse models significantly reduced the total neutrophil count in the lungs<sup>107</sup>. |
| MCL-1 | **Direct evidence;** MCL-1 function has been documented as a host factor manipulated by Mtb to promote bacterial survival<sup>108</sup>. The core strategy involves using small molecule inhibitors (BH3 mimetics) to neutralize MCL-1's anti-apoptotic function, thereby inducing intrinsic apoptosis in Mtb-infected macrophages<sup>109</sup>. |
| BAX | **Direct evidence;** BAX is the central apoptotic pathway against Mtb. Since BAX promotes apoptosis, its upregulation is directly associated with the therapeutic success of several proposed HDTs, opening important avenues for targeting apoptotic pathways<sup>109–112</sup>. |
| DDA1 | **Indirect evidence;** DDA1 directly regulates the NFκB signaling pathway, which is fundamental to host immune responses<sup>113</sup>. DDA1's strong correlation with an immune-suppressive microenvironment and its structural role as a component of a targetable ligase complex suggest it holds potential as a host-directed therapeutic target to reverse immune suppression or modulate chronic inflammation<sup>114</sup>. |
| SNAP23 | **Direct evidence;** Sources establish that SNAP-23 is a protein activator required for completing cargo secretion in secretory autophagy. This pathway is responsible for inflammatory responses. Literature suggests that inhibiting SNAP-23 function could be a strategy to block the unconventional secretion of pro-inflammatory cargos, thereby balancing protection against excessive pathology<sup>115</sup>. |
| ITGB1 | **Direct evidence;** ITGB1 is essential for forming VLA-4, which regulates the migration, accumulation, and in vivo function of a distinct, highly differentiated subset of T cells within mycobacterial lesions. Literature suggests leveraging the nitric oxide or IL-4 pathways to suppress ITGB1/VLA-4 expression to suppress granuloma development and control Mtb infection<sup>116</sup>. |
| ITGA5 | **Direct evidence;** Study suggests that ITGA5 acts as a novel receptor facilitating the entry of Mtb, as well as controlling downstream elements involved in apoptosis, oxidative stress response, and inflammatory signaling<sup>117</sup>. |
| MAP3PK4 | **Indirect evidence;** MAP3PK4 regulates p38 MAPK signaling, which controls inflammatory processes including pulmonary inflammation and the production of several key cytokines<sup>118</sup>. Suppression or inhibition of MAP3PK4 enriches T cells with stem-cell-like properties, targeting this kinase could be leveraged to generate a more persistent and robust T cell response, potentially enhancing host immunity against the persistent Mtb infection<sup>119</sup>. |
| MAP2K6 | **Direct evidence;** Literature indicates that MAP2K6 is a significant host factor responsible for cellular stress response, immune response, and inflammation regulation. Its inverse expression relationship with METTL14, linked through miR-29a-3p, offers the potential of METTL14 activators, miR-29a-3p mimics, or MAP2K6 inhibitors for TB therapeutics<sup>120</sup>. |

For instance, *IL-8* mediates immune response to *M. tuberculosis* by driving T cell recruitment, with evidence for a stage-dependent role beneficial early in infection^98–100^. *CXCR2* regulates neutrophil subpopulations and plays a critical role in TB immunopathogenesis. In TB-susceptible mouse models, selective CXCR2 inhibition significantly reduced pulmonary neutrophil counts^101^.

We selected drugs with known target genes forming the strongest connections (highest average betweenness centrality [BC]) to TB disease genes, narrowing to 23 high-confidence drugs from our 64 candidates (**Figure 4**). These include known TB HDTs, such as simvastatin, axitinib, resveratrol, and tamoxifen, and less-recognized candidates with high network proximity to TB disease genes.

Novel candidates include benzonatate (local anesthetic), bendroflumethiazide (sodium/potassium/chloride transporter inhibitor), amiodarone (potassium channel blocker), methylergometrine (dopamine receptor antagonist/serotonin receptor antagonist), lisuride (dopamine receptor agonist), chlorprothixene (dopamine receptor antagonist), methoxsalen (DNA synthesis inhibitor), and mitotane (antineoplastic agent) (**Figure 4**). Recovery of known TB-related genes among in-path genes supports the biological relevance of these novel predictions.

### 3.7 Synergistic MOA combinations for HDT candidates and TB antibiotics

Combining HDTs with pathogen-directed antibiotics can improve host response modulation by targeting multiple host-pathogen interaction pathways^3^. We investigated which HDT candidates and TB antibiotics function synergistically. For each drug pair, we computed synergy scores using four metrics and mapped results to corresponding MOAs to summarize synergy across all MOA pairs (**Methods 2.10**).

Our analysis predicted potentially complementary HDT MOAs to TB antibiotics (**Figure 5**). Antineoplastic agents showed promising synergy with bacterial cell wall synthesis inhibitors. While not evaluated in TB, their potential for combinatorial synergy is corroborated by *in vitro* evidence in cancer^121^. This supports a possible mechanistic basis for the predicted synergy, motivating experimental evaluation of antineoplastic agents and bacterial cell wall synthesis inhibitors as an adjunct host-directed strategy in TB.

**Figure 5.**
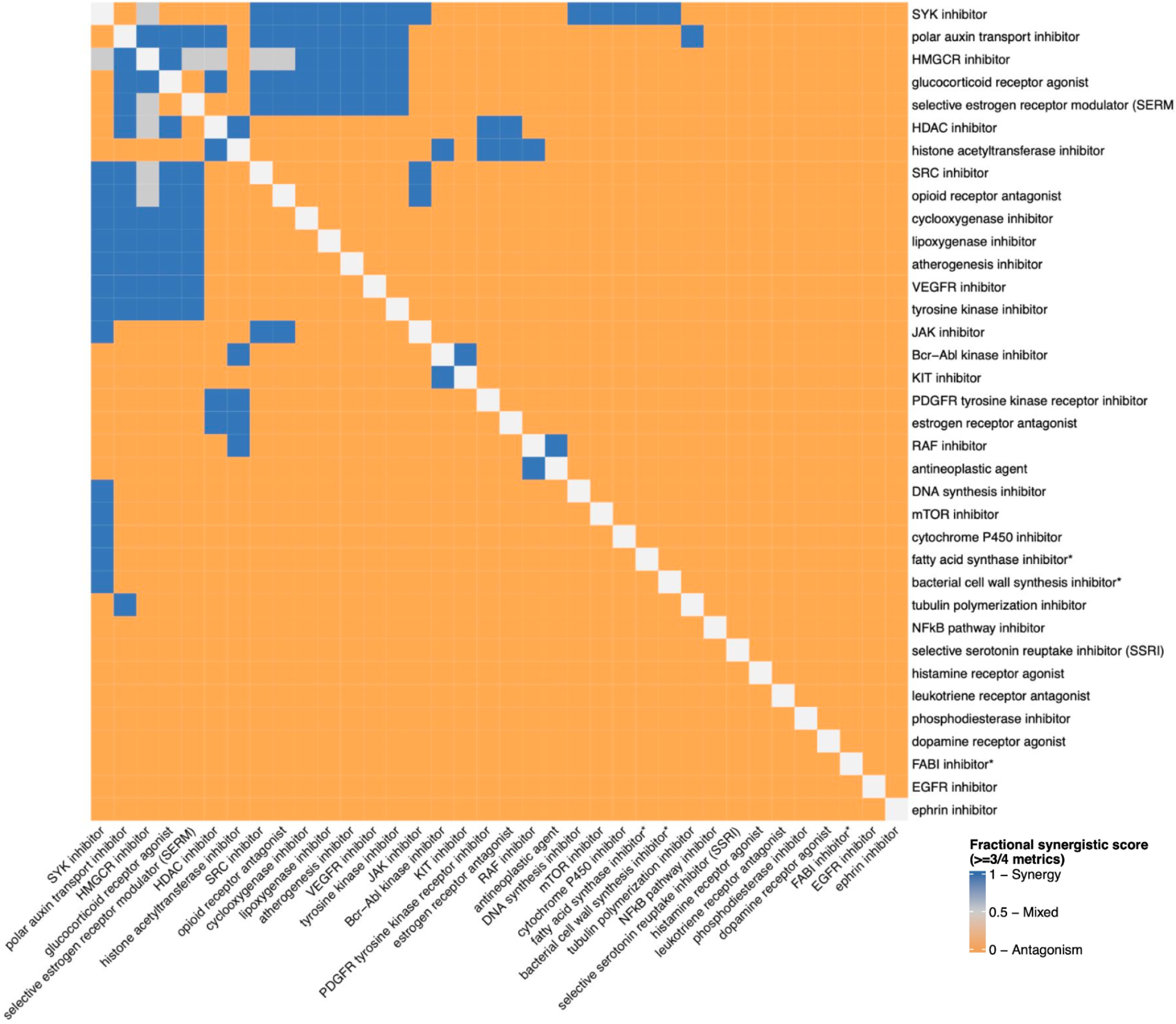
Synergy and antagonism of pairs of mechanisms of action (MOA) associated with our 64 high-confidence drug candidates and current TB antibiotics. Heatmap shows synergy voting fraction for all pairwise MOA combinations from our HDT candidates and TB antibiotics (noted with asterisk [*]). Synergy scores were computed using the four metrics (ZIP, Bliss, Loewe, and HSA) from *DrugCombDB*. Drug pairs were labeled synergy (or antagonism) if ≥3/4 DrugCombDB metrics agreed, and the fractional synergistic score for each MOA pair was computed as the proportion of its drug pairs classified as synergistic, counting each drug pair once (blue = predicted synergy; orange = predicted antagonism; grey = inconclusive; white = no values for self-pairs).

We also predicted synergy between bacterial cell wall synthesis inhibitors and tubulin polymerization inhibitors. This is biologically feasible because bacterial cell division and cell wall construction are closely linked, and some tubulin-targeting drugs limit *M. tuberculosis* growth^122–124^. Additionally, altering microtubule function can affect immune cell control of intracellular bacteria, as suggested by host-directed activity of colchicine^125^. While direct TB combination studies are limited, these findings support the predicted interaction. Further, we predicted antagonism between bacterial cell-wall synthesis inhibitors and fatty acid synthesis inhibitors because fatty-acid synthesis inhibitors slow bacterial growth. Cell-wall antibiotics work best when bacteria actively build new cell walls, so slowing this process reduces cell-wall drugs’ effectiveness when combined^126^.

Among synergistic HDT MOA pairs (**Figure 5**), we observed literature-supported synergy between HMGCR inhibitors (e.g., statins) and HDAC inhibitors (e.g., resveratrol), which show cooperative effects on autophagy induction and immune modulation, suggesting enhanced host antimicrobial responses during TB infection^127^. Predicted synergy between KIT pathway inhibitors and BCR-ABL kinase inhibitors is supported by multi-target inhibitors such as dasatinib, which simultaneously inhibits both pathways. This demonstrates that co-targeting KIT and BCR-ABL signaling is pharmacologically feasible and produces coordinated suppression of shared downstream growth and survival programs^128,129^.

Our analysis identified several antagonistic MOA combinations, particularly those involving receptor-modulating or ion channel-targeting pathways, including selective estrogen receptor modulators, dopamine receptor agonists, and opioid receptor antagonists. While these pairs showed negative synergy scores, we found no evidence that these MOAs are mutually antagonistic.

This analysis provides a guide for prioritizing synergistic drug mechanism combinations and avoiding antagonistic interactions (especially with comorbidity treatments) for both HDT-antibiotic combinations and HDT-HDT combinations in TB.

## 4. Discussion

TB urgently needs new host-directed therapeutics that can complement existing antibiotics and address rising drug resistance. Transcriptome-based drug repurposing offers a promising approach by identifying FDA-approved drugs that reverse disease gene expression patterns. However, current TB repurposing efforts face significant challenges: disease signatures vary across profiling technologies, biological contexts, and analytical pipelines, while most studies rely on single connectivity methods and limited datasets^20–22^, which can bias drug predictions and miss robust candidates^23^.

We developed an integrative computational framework to address these limitations. Our approach combines 28 TB gene expression datasets across multiple platforms, biological contexts, and cell types with six complementary connectivity scoring methods. Critically, we introduce a weighted aggregation strategy to derive consensus disease signatures that capture dominant host responses while mitigating dataset-specific biases.

### 4.1 Summary of key findings

Applying this approach, we successfully identified 64 high-confidence candidate HDTs for TB, including both known drugs and novel predictions **(Table 1; Table S2**). As proof-of-principle, our workflow successfully prioritized known TB HDTs, including statins and the experimentally validated candidate tamoxifen, validating our integrative approach. Among novel predictions, clonidine, an α2-adrenergic receptor agonist used for hypertension, emerged as a candidate with documented effects on immune and inflammatory responses^130–132^ suggesting potential for modulating TB immunopathology. These predictions demonstrate that our framework recovers established potential HDTs while identifying therapeutically plausible new candidates.

Top predicted HDTs were enriched for diverse host-targeted mechanisms including HMGCR inhibition, PDGFR tyrosine kinase inhibition, adrenergic receptor antagonism, cyclooxygenase inhibition, and estrogen receptor modulation (**Table 2**). These mechanisms target complementary aspects of TB pathogenesis, e.g., lipid metabolism, growth-factor signaling, inflammatory pathways, and hormonal regulation, that influence macrophage function, immune responses, and infection-associated pathology, providing multiple routes for host-directed intervention.

### 4.2 Advancing transcriptome-based drug repurposing for infectious diseases

Integrating heterogeneous TB transcriptomic datasets and multiple connectivity metrics substantially altered drug prioritization compared to prior connectivity-based approaches. Rather than relying on a single disease-drug reversal score or narrowly defined disease signature, our framework emphasized candidates whose transcriptional effects aligned with dominant host responses across diverse TB contexts. This integration reduced sensitivity to dataset-specific biases and enabled recovery of HDT candidates robust across tissues, platforms, and experimental settings.

By prioritizing drugs that reverse both aggregated and individual TB signatures, our approach captures conserved biological processes and identifies candidates that shift host responses toward a healthier state. Recovery of known HDTs (e.g., statins, tamoxifen) alongside novel candidates validates our integrative workflow for TB drug repurposing and demonstrates its potential for host-directed therapy development in other infectious diseases.

### 4.3 Methodological innovations and their impact

Transcriptomic responses to TB infection vary substantially across profiling technologies and biological conditions. Our workflow addresses this heterogeneity through weighted aggregation of multiple disease signatures. Our strategy effectively captures dominant disease signals, core host-response pathways observed across the individual signatures (**Figure 3**). This is evidenced by pathway retention and drug overlap; among 588 drugs identified using individual signatures, 214 (36%) were recovered in aggregated signatures, representing candidates that consistently reversed TB signatures across datasets. This overlap supports that aggregation preserves biologically meaningful signals while filtering dataset-specific noise.

Variance analysis revealed greater variability in GO:BP enrichment for blood-only aggregated signatures compared to aggregation across tissue types (blood, BAL, and cell lines; **Figure S3; Methods 2.7.2**). This demonstrates that heterogeneity persists even within similar biological contexts. Therefore, we recommend incorporating diverse signatures during aggregation rather than restricting to closely related contexts. Aggregating across platforms, tissues, and experimental conditions favors pathways consistently enriched across diverse settings and reduces bias from any single source, enabling identification of robust therapeutic candidates despite underlying dataset heterogeneity.

In addition to signature aggregation, we implemented six complementary methods to quantify drug reversal: four enrichment-based (CMAP 1.0, CMAP 2.0: WCS, NCS, τ) and two correlation-based (XCor, XSpe). These methods capture distinct aspects of disease-drug relationships and introduce method-specific biases that impact drug rankings (**Table S2**). We applied rank-based aggregation across scoring modalities to prioritize drugs consistently highly ranked across methods, increasing stability and interpretability. This multi-method integration successfully recovered known TB HDT candidates (statins, tamoxifen) with consistent high rankings across methods (**Tables 1; S2**), validating our approach.

Beyond global reversal patterns, network analysis provided mechanistic insights by identifying 12 key linking genes (including *IL-8*, *CXCR2*) that bridge TB disease genes and drug targets within the protein-protein interaction network (**Figure 4**; **Table 3**). This analysis also revealed convergence of predicted HDTs on cholesterol and cytokine pathways through the shared node HMGCS1 (**Figure S7**). Additionally, our MOA-level synergy analysis identified pharmacologically feasible combinations, including co-targeting KIT and BCR-ABL signaling, supporting multi-target kinase inhibitors like dasatinib (**Figure 5**).

### 4.4 Limitations and technical considerations

The LINCS drug perturbation dataset exhibits variation across cell lines, dosages, and treatment durations (**Methods 2.4**). For consistency, we restricted analysis to 10 μM concentration with a 24-hour treatment, the most common drug condition, representing 27% of LINCS signatures^41,42^, and summarized drug effects in a cell line-agnostic manner (**Methods 2.6**). Limited cell type diversity in LINCS remains a constraint. We evaluated biological matches between disease control samples and untreated LINCS cell lines by correlation analysis (**Methods 2.11**). While some tissue-specific structure emerged, drug-disease correlations were consistently weak whereas drug-drug correlations were strong (**Figure S9**). This indicates that baseline-matching by direct correlation is inadequate for disease-drug comparison and improved methods are needed to define biologically meaningful baselines for connectivity mapping.

We restricted disease-drug reversal scoring to the 978 L1000 genes^12^, a representative set whose directly measured expression enables robust transcriptome inference. This constraint improves consistency and avoids noise from inferred values, but reduces gene coverage and may exclude disease-associated genes absent in the L1000 set. Future work could incorporate inferred genes with confidence weighting or validate findings using transcriptome-wide data sources, such as CycleGAN-based drug signatures predictions^133^. Ultimately, experimental validation is necessary to confirm therapeutic efficacy of prioritized TB HDT candidates, both as single agents and in synergistic combinations with TB antibiotics or HDTs.

### 4.5 Therapeutic implications and future directions

Our approach successfully identified known and novel HDT candidates for TB, demonstrating potential for HDT repurposing in other infectious diseases. MOA-level synergy analysis enables rational design of combination therapies with pathogen-directed antibiotics by identifying drug pairs with complementary or convergent mechanisms that may act synergistically.

Literature evidence corroborates predicted synergistic combinations. HMGCR inhibitors (statins) combined with HDAC inhibitors (e.g., resveratrol) may synergistically induce autophagy and modulate immune responses during TB infection^127^. Multi-target kinase inhibitors like dasatinib, which simultaneously inhibit KIT and BCR-ABL pathways^128,129^, demonstrate pharmacological feasibility of co-targeting these signaling pathways. These pathways converge on downstream programs controlling cell survival and proliferation, supporting predicted synergy between inhibitors of KIT receptor tyrosine kinase and BCR-ABL kinase^134^ (**Figure 5**).

We also predicted antagonistic MOA combinations, though mechanistic evidence is limited. For example, fatty acid synthesis inhibitors may antagonize bacterial cell wall synthesis inhibitors because slowing bacterial growth reduces cell wall turnover. Cell wall-active antibiotics work best against dividing bacteria, so reduced growth may diminish their joint efficacy^126,135,136^. These predicted antagonisms warrant experimental validation to determine clinical relevance.

Finally, network-based analysis revealed 12 novel druggable targets (including *IL-8*, *CXCR2*) that mediate interactions between TB disease genes and drug targets within the protein-protein interaction network (**Figure 4**; **Table 3**). This systems-level understanding reinforces the rationale for targeting specific host pathways and supports combinatorial therapeutic strategies. *In vitro* and *in vivo* validation of prioritized candidates, as single agents, and in combination with TB antibiotics, is essential to assess therapeutic efficacy. Future work should incorporate drug cell-line matching to identify optimal contexts for experimental validation of HDTs and their targets. This integrative framework provides a generalizable approach for HDT discovery for tuberculosis and other infectious diseases.

## 5. Conclusion

In summary, we developed and validated an integrative computational framework that addresses fundamental challenges in heterogeneous transcriptome-based drug repurposing for tuberculosis. While heterogeneity in biological and technical conditions across datasets remains a challenge, our approach is specifically designed to prioritize robust HDT candidate predictions despite such variability. By combining weighted signature aggregation with multiple connectivity methods and network-based analysis, we identified 64 high-confidence HDT candidates and 12 novel druggable targets. This work establishes a robust, generalizable approach for systematic host-directed therapeutic discovery for infectious diseases.

## Data and code availability

All processed disease data and code used to perform the analyses and generate the results and figures presented in this study are publicly available at github.com/JRaviLab/integrative-drugrep-tb.

## Author contributions

KS obtained the microarray data and constructed the associated disease signatures, designed and built the workflow, performed disease-drug comparison and other downstream analyses, acquired and interpreted results, and drafted and revised the manuscript. LT updated microarray and RNAseq datasets and revised the codebase. LB conducted the baseline comparison analysis. AT obtained the RNAseq data and constructed RNAseq disease signatures. JR and AK conceived the study, provided guidance throughout the project, and contributed to the design of the workflow, interpretation of the results, and revision of the manuscript.

## Acknowledgment

We thank Keenan Manpearl for providing propagated annotations of GO:BP terms used in our variable pathway analyses. Special thanks to Dr. Christina Stallings, the Washington University St. Louis and Emily Meyer from the Department of Biomedical Informatics, CU Anschutz, for their insightful feedback on the manuscript. We also would like to thank the current and past members of the JRavi and Krishnan labs for support, especially Evan Brenner and Abhirupa Ghosh for insightful feedback and discussions, and Emily Boyer and Jacob Krol, for their valuable feedback on the manuscript.

## References

1. World Health Organization. Global Tuberculosis Report 2023. (2023).

2. Dheda, K. et al. The epidemiology, pathogenesis, transmission, diagnosis, and management of multidrug-resistant, extensively drug-resistant, and incurable tuberculosis. Lancet Respir. Med. 5, 291–360 (2017).

3. Kaufmann, S. H. E., Dorhoi, A., Hotchkiss, R. S. & Bartenschlager, R. Host-directed therapies for bacterial and viral infections. Nat. Rev. Drug Discov. 17, 35–56 (2018).

4. Machelart, A., Song, O.-R., Hoffmann, E. & Brodin, P. Host-directed therapies offer novel opportunities for the fight against tuberculosis. Drug Discov. Today 22, 1250–1257 (2017).

5. Kilinç, G., Saris, A., Ottenhoff, T. H. M. & Haks, M. C. Host-directed therapy to combat mycobacterial infections*. Immunol. Rev. 301, 62–83 (2021).

6. Wallis, R. S. et al. Tuberculosis--advances in development of new drugs, treatment regimens, host-directed therapies, and biomarkers. Lancet Infect. Dis. 16, e34–46 (2016).

7. Zumla, A. et al. Towards host-directed therapies for tuberculosis. Nat. Rev. Drug Discov. 14, 511–512 (2015).

8. Hawn, T. R., Matheson, A. I., Maley, S. N. & Vandal, O. Host-Directed Therapeutics for Tuberculosis: Can We Harness the Host? Microbiol. Mol. Biol. Rev. 77, 608–627 (2013).

9. Karaman, B. & Sippl, W. Computational Drug Repurposing: Current Trends. Curr. Med. Chem. 26, 5389–5409 (2019).

10. Cousins, H. C., Nayar, G. & Altman, R. B. Computational Approaches to Drug Repurposing: Methods, Challenges, and Opportunities. Annu. Rev. Biomed. Data Sci. 7, 15–29 (2024).

11. Lamb, J. The Connectivity Map: a new tool for biomedical research. Nat. Rev. Cancer 7, 54–60 (2007).

12. Subramanian, A. et al. A Next Generation Connectivity Map: L1000 Platform and the First 1,000,000 Profiles. Cell 171, 1437–1452.e17 (2017).

13. Samart, K., Tuyishime, P., Krishnan, A. & Ravi, J. Reconciling multiple connectivity scores for drug repurposing. Brief. Bioinform. bbab161 (2021) doi:10.1093/bib/bbab161.

14. Atri, P. et al. Connectivity mapping-based identification of pharmacological inhibitor targeting HDAC6 in aggressive pancreatic ductal adenocarcinoma. *Npj Precis*. Oncol. 8, 66 (2024).

15. Raghavan, R. et al. Drug discovery using clinical outcome-based Connectivity Mapping: application to ovarian cancer. BMC Genomics 17, 811 (2016).

16. Zhao, Y., Chen, X., Chen, J. & Qi, X. Decoding Connectivity Map-based drug repurposing for oncotherapy. Brief. Bioinform. 24, bbad142 (2023).

17. Piastra, V. et al. Repurposed AT9283 triggers anti-tumoral effects by targeting MKK3 oncogenic functions in Colorectal Cancer. J. Exp. Clin. Cancer Res. 43, 234 (2024).

18. Cummings, J. L. et al. Drug repurposing for Alzheimer’s disease and other neurodegenerative disorders. Nat. Commun. 16, 1755 (2025).

19. Williams, G. et al. Drug repurposing for Alzheimer’s disease based on transcriptional profiling of human iPSC-derived cortical neurons. Transl. Psychiatry 9, 220 (2019).

20. Wang, Z., Arat, S., Magid-Slav, M. & Brown, J. R. Meta-analysis of human gene expression in response to Mycobacterium tuberculosis infection reveals potential therapeutic targets. BMC Syst. Biol. 12, 3 (2018).

21. Jiang, Y., Zhang, J.-X. & Liu, R. Systematic comparison of differential expression networks in MTB mono-, HIV mono- and MTB/HIV co-infections for drug repurposing. PLOS Comput. Biol. 18, e1010744 (2022).

22. Deng, S. et al. Integrated bioinformatic analyses investigate macrophage-M1-related biomarkers and tuberculosis therapeutic drugs. Front. Genet. 14, 1041892 (2023).

23. Lim, N. & Pavlidis, P. Evaluation of connectivity map shows limited reproducibility in drug repositioning. Sci. Rep. 11, 17624 (2021).

24. Gonzalez Gomez, C., Rosa-Calatrava, M. & Fouret, J. Optimizing *in silico* drug discovery: simulation of connected differential expression signatures and applications to benchmarking. Brief. Bioinform. 25, bbae299 (2024).

25. Cheng, J., Yang, L., Kumar, V. & Agarwal, P. Systematic evaluation of connectivity map for disease indications. Genome Med. 6, 540 (2014).

26. Cheng, J. & Yang, L. Comparing gene expression similarity metrics for connectivity map. in 2013 IEEE International Conference on Bioinformatics and Biomedicine 165–170 (IEEE, Shanghai, China, 2013). doi:10.1109/BIBM.2013.6732481.

27. Paton, V. et al. Assessing the impact of transcriptomics data analysis pipelines on downstream functional enrichment results. Nucleic Acids Res. 52, 8100–8111 (2024).

28. Wang, B., Van Der Kloet, F., Kes, M. B. M. J., Luirink, J. & Hamoen, L. W. Improving gene set enrichment analysis (GSEA) by using regulation directionality. Microbiol. Spectr. 12, e03456–23 (2024).

29. Di Iulio, J., Bartha, I., Spreafico, R., Virgin, H. W. & Telenti, A. Transfer transcriptomic signatures for infectious diseases. Proc. Natl. Acad. Sci. 118, e2022486118 (2021).

30. Dutta, N. K. et al. Statin adjunctive therapy shortens the duration of TB treatment in mice. J. Antimicrob. Chemother. 71, 1570–1577 (2016).

31. Alffenaar, J.-W. C., Akkerman, O. W. & van Hest, R. Statin Adjunctive Therapy for Tuberculosis Treatment. Antimicrob. Agents Chemother. 60, 7004 (2016).

32. Guerra-De-Blas, P. D. C. et al. Potential Effect of Statins on Mycobacterium tuberculosis Infection. J. Immunol. Res. 2018, 7617023 (2018).

33. Guerra-De-Blas, P. D. C. et al. Simvastatin Enhances the Immune Response Against Mycobacterium tuberculosis. Front. Microbiol. 10, 2097 (2019).

34. Boland, R. et al. Repurposing Tamoxifen as Potential Host-Directed Therapeutic for Tuberculosis. mBio 14, e03024–22 (2023).

35. Clough, T., E. ,. &. Barrett. The Gene Expression Omnibus Database. 10.1007/978-1-4939-3578-9_5 (2016) 10.1007/978-1-4939-3578-9_5.

36. Greene, C. S., et al. refine.bio: a resource of uniformly processed publicly available gene expression datasets. (2023).

37. Lachmann, A. et al. Massive mining of publicly available RNA-seq data from human and mouse. Nat. Commun. 9, 1366 (2018).

38. Ritchie, M. E. et al. limma powers differential expression analyses for RNA-sequencing and microarray studies. Nucleic Acids Res. 43, e47 (2015).

39. Love, M. I., Huber, W. & Anders, S. Moderated estimation of fold change and dispersion for RNA-seq data with DESeq2. Genome Biol. 15, 550 (2014).

40. Duan, Y. et al. signatureSearch: environment for gene expression signature searching and functional interpretation. Nucleic Acids Res. 48, e124 (2020).

41. Chen, B. et al. Reversal of cancer gene expression correlates with drug efficacy and reveals therapeutic targets. Nat. Commun. 8, 16022 (2017).

42. Tong, X. et al. Deep representation learning of chemical-induced transcriptional profile for phenotype-based drug discovery. Nat. Commun. 15, 5378 (2024).

43. Corsello, S. M. et al. The Drug Repurposing Hub: a next-generation drug library and information resource. Nat. Med. 23, 405–408 (2017).

44. Zhou, X., Wang, M., Katsyv, I., Irie, H. & Zhang, B. EMUDRA: Ensemble of Multiple Drug Repositioning Approaches to improve prediction accuracy. Bioinforma. Oxf. Engl. 34, 3151–3159 (2018).

45. Zhang, S.-D. & Gant, T. W. A simple and robust method for connecting small-molecule drugs using gene-expression signatures. BMC Bioinformatics 9, 258 (2008).

46. A, S., et al. Gene set enrichment analysis: a knowledge-based approach for interpreting genome-wide expression profiles. Proc. Natl. Acad. Sci. U. S. A. 102, (2005).

47. Spearman Rank Correlation Coefficient. in The Concise Encyclopedia of Statistics 502–505 (Springer New York, New York, NY, 2008). doi:10.1007/978-0-387-32833-1_379.

48. Wang, B. et al. Systematic comparison of ranking aggregation methods for gene lists in experimental results. Bioinformatics 38, 4927–4933 (2022).

49. Li, X., Choudhary, P. K., Biswas, S. & Wang, X. A Bayesian latent variable approach to aggregation of partial and top-ranked lists in genomic studies. Stat. Med. 37, 4266–4278 (2018).

50. Ashburner, M. et al. Gene Ontology: tool for the unification of biology. Nat. Genet. 25, 25–29 (2000).

51. Wu, T. et al. clusterProfiler 4.0: A universal enrichment tool for interpreting omics data. The Innovation 2, 100141 (2021).

52. Fröhlich, H., Speer, N., Poustka, A. & Beißbarth, T. GOSim – an R-package for computation of information theoretic GO similarities between terms and gene products. BMC Bioinformatics 8, 166 (2007).

53. Cooper, A. M., Mayer-Barber, K. D. & Sher, A. Role of innate cytokines in mycobacterial infection. Mucosal Immunol. 4, 252–260 (2011).

54. Cannon, M. et al. DGIdb 5.0: rebuilding the drug–gene interaction database for precision medicine and drug discovery platforms. Nucleic Acids Res. 52, D1227–D1235 (2024).

55. Szklarczyk, D. et al. The STRING database in 2023: protein–protein association networks and functional enrichment analyses for any sequenced genome of interest. Nucleic Acids Res. 51, D638–D646 (2023).

56. Dijkstra, E. W. A note on two problems in connexion with graphs. Numer. Math. 1, 269–271 (1959).

57. Padda, I. S. & Muralidhara Reddy, K. Antitubercular Medications. in StatPearls (StatPearls Publishing, Treasure Island (FL), 2025).

58. Liu, H. et al. DrugCombDB: a comprehensive database of drug combinations toward the discovery of combinatorial therapy. Nucleic Acids Res. gkz1007 (2019) doi:10.1093/nar/gkz1007.

59. Yadav, B., Wennerberg, K., Aittokallio, T. & Tang, J. Searching for Drug Synergy in Complex Dose-Response Landscapes Using an Interaction Potency Model. Comput. Struct. Biotechnol. J. 13, 504–513 (2015).

60. Bliss, C. I. THE TOXICITY OF POISONS APPLIED JOINTLY1. Ann. Appl. Biol. 26, 585–615 (1939).

61. Loewe, S. Die quantitativen Probleme der Pharmakologie. Ergeb. Physiol. 27, 47–187 (1928).

62. Berenbaum, M. C. What is synergy? Pharmacol. Rev. 41, 93–141 (1989).

63. Webber, W., Moffat, A. & Zobel, J. A similarity measure for indefinite rankings. ACM Trans. Inf. Syst. 28, 1–38 (2010).

64. Novikov, A., et al. *Mycobacterium tuberculosis* Triggers Host Type I IFN Signaling To Regulate IL-1β Production in Human Macrophages. J. Immunol. 187, 2540–2547 (2011).

65. Berry, M. P. R. et al. An interferon-inducible neutrophil-driven blood transcriptional signature in human tuberculosis. Nature 466, 973–977 (2010).

66. Raju, B. et al. Gene expression profiles of bronchoalveolar cells in pulmonary TB. Tuberculosis 88, 39–51 (2008).

67. Akter, S. et al. Mycobacterium tuberculosis infection drives a type I IFN signature in lung lymphocytes. Cell Rep. 39, 110983 (2022).

68. Andreu, N. et al. Primary macrophages and J774 cells respond differently to infection with Mycobacterium tuberculosis. Sci. Rep. 7, 42225 (2017).

69. Shenoy, A. T. et al. Antigen presentation by lung epithelial cells directs CD4+ TRM cell function and regulates barrier immunity. Nat. Commun. 12, 5834 (2021).

70. Bachanová, P. et al. Comparative transcriptomic analysis of whole blood mycobacterial growth assays and tuberculosis patients’ blood RNA profiles. Sci. Rep. 12, 17684 (2022).

71. Gao, X. et al. An updated comparison of microarray and RNA-seq for concentration response transcriptomic study: case studies with two cannabinoids, cannabichromene and cannabinol. BMC Genomics 26, 392 (2025).

72. Rai, M. F., Tycksen, E. D., Sandell, L. J. & Brophy, R. H. Advantages of RNA-seq compared to RNA microarrays for transcriptome profiling of anterior cruciate ligament tears. J. Orthop. Res. 36, 484–497 (2018).

73. Zhang, Y., Lin, Y., Yu, H., Tian, R. & Li, F. Characteristic genes in THP-1 derived macrophages infected with Mycobacterium tuberculosis H37Rv strain identified by integrating bioinformatics methods. Int. J. Mol. Med. 10.3892/ijmm.2019.4293 (2019) doi:10.3892/ijmm.2019.4293.

74. Fontán, P., Aris, V., Ghanny, S., Soteropoulos, P. & Smith, I. Global Transcriptional Profile of *Mycobacterium tuberculosis* during THP-1 Human Macrophage Infection. Infect. Immun. 76, 717–725 (2008).

75. Lopes-Ramos, C. M. et al. Sex Differences in Gene Expression and Regulatory Networks across 29 Human Tissues. Cell Rep. 31, 107795 (2020).

76. Dutta, N. K. et al. Adjunctive Host-Directed Therapy With Statins Improves Tuberculosis-Related Outcomes in Mice. J. Infect. Dis. 221, 1079–1087 (2020).

77. Adewole, O. O. et al. Atorvastatin accelerates *Mycobacterium tuberculosis* clearance in pulmonary TB: a randomised phase IIA trial. Int. J. Tuberc. Lung Dis. 27, 226–228 (2023).

78. Davuluri, K. S. et al. Atorvastatin Potentially Reduces Mycobacterial Severity through Its Action on Lipoarabinomannan and Drug Permeability in Granulomas. Microbiol. Spectr. 11, e03197–22 (2023).

79. Montero-Vega, M. T. et al. Fluvastatin Converts Human Macrophages into Foam Cells with Increased Inflammatory Response to Inactivated Mycobacterium tuberculosis H37Ra. Cells 13, 536 (2024).

80. Jhilta, A. et al. Host-Directed Therapy with Inhalable Lovastatin Microspheres for Matrix Metalloproteinase Inhibition in Tuberculosis. ACS Appl. Bio Mater. 8, 1533–1546 (2025).

81. Cross, G. B. et al. Rosuvastatin adjunctive therapy for rifampicin-susceptible pulmonary tuberculosis: a phase 2b, randomised, open-label, multicentre trial. Lancet Infect. Dis. 23, 847–855 (2023).

82. Cross, G. B. et al. PET-CT outcomes from a randomised controlled trial of rosuvastatin as an adjunct to standard tuberculosis treatment. Nat. Commun. 15, 10475 (2024).

83. Mourenza, Á., Gil, J. A., Mateos, L. M. & Letek, M. Novel Treatments against Mycobacterium tuberculosis Based on Drug Repurposing. Antibiot. Basel Switz. 9, 550 (2020).

84. Maiga, M. et al. Efficacy of Adjunctive Tofacitinib Therapy in Mouse Models of Tuberculosis. EBioMedicine 2, 868–873 (2015).

85. Mahon, R. N. & Hafner, R. Immune Cell Regulatory Pathways Unexplored as Host-Directed Therapeutic Targets for *Mycobacterium tuberculosis* : An Opportunity to Apply Precision Medicine Innovations to Infectious Diseases. Clin. Infect. Dis. 61, S200–S216 (2015).

86. Chugh, S., et al. Polyphosphate kinase-1 regulates bacterial and host metabolic pathways involved in pathogenesis of *Mycobacterium tuberculosis*. Proc. Natl. Acad. Sci. 121, e2309664121 (2024).

87. Wehrstedt, S. et al. The tyrosine kinase inhibitor dasatinib reduces the growth of intracellular *Mycobacterium tuberculosis* despite impairing T-cell function. Eur. J. Immunol. 48, 1892–1903 (2018).

88. Shankaran, D. et al. The antidepressant sertraline provides a novel host directed therapy module for augmenting TB therapy. eLife 12, e64834 (2023).

89. Young, C., Walzl, G. & Du Plessis, N. Therapeutic host-directed strategies to improve outcome in tuberculosis. Mucosal Immunol. 13, 190–204 (2020).

90. Korbee, C. J. et al. Combined chemical genetics and data-driven bioinformatics approach identifies receptor tyrosine kinase inhibitors as host-directed antimicrobials. Nat. Commun. 9, 358 (2018).

91. Napier, R. J. et al. Imatinib-Sensitive Tyrosine Kinases Regulate Mycobacterial Pathogenesis and Represent Therapeutic Targets against Tuberculosis. Cell Host Microbe 10, 475–485 (2011).

92. Chai, Q. et al. A Mycobacterium tuberculosis surface protein recruits ubiquitin to trigger host xenophagy. Nat. Commun. 10, 1973 (2019).

93. Mortensen, R. et al. Cyclooxygenase inhibitors impair CD4 T cell immunity and exacerbate Mycobacterium tuberculosis infection in aerosol-challenged mice. *Commun*. Biol. 2, 288 (2019).

94. Vilaplana, C. et al. Ibuprofen Therapy Resulted in Significantly Decreased Tissue Bacillary Loads and Increased Survival in a New Murine Experimental Model of Active Tuberculosis. J. Infect. Dis. 208, 199–202 (2013).

95. Nore, K. G. et al. The Cyclooxygenase 2 Inhibitor Etoricoxib as Adjunctive Therapy in Tuberculosis Impairs Macrophage Control of Mycobacterial Growth. J. Infect. Dis. 229, 888–897 (2024).

96. Gan, Y., Hu, Q., Li, A., Gu, L. & Guo, S. Estradiol inhibits autophagy of *Mycobacterium tuberculosis* -infected 16HBE cells and controls the proliferation of intracellular *Mycobacterium tuberculosis*. Mol. Med. Rep. 25, 196 (2022).

97. Gupta, M., Srikrishna, G., Klein, S. L. & Bishai, W. R. Genetic and hormonal mechanisms underlying sex-specific immune responses in tuberculosis. Trends Immunol. 43, 640–656 (2022).

98. Wang, L. et al. Olink proteomics and lipidomics analysis of serum from patients infected with non-tuberculous mycobacteria and Mycobacterium tuberculosis. Inflamm. Res. 73, 1945–1960 (2024).

99. Krupa, A. et al. Binding of CXCL8/IL-8 to *Mycobacterium tuberculosis* Modulates the Innate Immune Response. Mediators Inflamm. 2015, 124762 (2015).

100. Diatlova, A. et al. Molecular Markers of Early Immune Response in Tuberculosis: Prospects of Application in Predictive Medicine. Int. J. Mol. Sci. 24, 13261 (2023).

101. Saqib, M. et al. Pathogenic role for CD101-negative neutrophils in the type I interferon-mediated immunopathogenesis of tuberculosis. Cell Rep. 44, 115072 (2025).

102. Alfarsi, L. H. et al. PPFIA1 expression associates with poor response to endocrine treatment in luminal breast cancer. BMC Cancer 20, 425 (2020).

103. Gu, C. et al. Discovery of the Oncogenic Parp1, a Target of bcr-abl and a Potential Therapeutic, in mir-181a/PPFIA1 Signaling Pathway. Mol. Ther. - Nucleic Acids 16, 1–14 (2019).

104. Su, R. et al. PPFIA1-targeting miR-181a mimic and saRNA overcome imatinib resistance in BCR-ABL1-independent chronic myeloid leukemia by suppressing leukemia stem cell regeneration. Mol. Ther. - Nucleic Acids 32, 729–742 (2023).

105. Yuan, H. et al. Lysine catabolism reprograms tumour immunity through histone crotonylation. Nature 617, 818–826 (2023).

106. Mohammadnabi, N., et al. *Mycobacterium tuberculosis* : The Mechanism of Pathogenicity, Immune Responses, and Diagnostic Challenges. J. Clin. Lab. Anal. 38, e25122 (2024).

107. Alaridah, N. et al. Mycobacteria Manipulate G-Protein-Coupled Receptors to Increase Mucosal Rac1 Expression in the Lungs. J. Innate Immun. 9, 318–329 (2017).

108. Arnett, E. et al. Combination of MCL-1 and BCL-2 inhibitors is a promising approach for a host-directed therapy for tuberculosis. Biomed. Pharmacother. 168, 115738 (2023).

109. Arnett, E. et al. PPARγ is critical for Mycobacterium tuberculosis induction of Mcl-1 and limitation of human macrophage apoptosis. PLOS Pathog. 14, e1007100 (2018).

110. Bao, J. et al. Inhibition of mycobacteria proliferation in macrophages by low cisplatin concentration through phosphorylated p53-related apoptosis pathway. PLOS ONE 18, e0281170 (2023).

111. Zhang, Y., et al. *DHX36* , *BAX* , and *ARPC1B* May Be Critical for the Diagnosis and Treatment of Tuberculosis. Can. Respir. J. 2020, 1–11 (2020).

112. Upadhyay, R. et al. Host Directed Therapy for Chronic Tuberculosis via Intrapulmonary Delivery of Aerosolized Peptide Inhibitors Targeting the IL-10-STAT3 Pathway. Sci. Rep. 8, 16610 (2018).

113. Zhao, S. et al. DDA1 promotes stage IIB-IIC colon cancer progression by activating NFκB/CSN2/GSK-3β signaling. Oncotarget 7, 19794–19812 (2016).

114. Wu, F., Ning, H., Sun, Y., Wu, H. & Lyu, J. Integrative exploration of the mutual gene signatures and immune microenvironment between benign prostate hyperplasia and castration-resistant prostate cancer. Aging Male 26, 2183947 (2023).

115. Kimura, T. et al. Dedicated SNARE s and specialized TRIM cargo receptors mediate secretory autophagy. EMBO J. 36, 42–60 (2017).

116. Pearl, J. E. et al. Nitric oxide inhibits the accumulation of CD 4^+^ CD 44^hi^ T bet^+^ CD 69^lo^ T cells in mycobacterial infection. Eur. J. Immunol. 42, 3267–3279 (2012).

117. Zhang, J. Transcriptome Analysis Reveals Novel Entry Mechanisms and a Central Role of SRC in Host Defense during High Multiplicity Mycobacterial Infection. PLoS ONE 8, e65128 (2013).

118. Gupta, A., Ökesli-Armlovich, A., Morgens, D., Bassik, M. C. & Khosla, C. A genome-wide analysis of targets of macrolide antibiotics in mammalian cells. J. Biol. Chem. 295, 2057–2067 (2020).

119. Song, F. et al. A multi-kinase inhibitor screen identifies inhibitors preserving stem-cell-like chimeric antigen receptor T cells. Nat. Immunol. 26, 279–293 (2025).

120. Feng, L., Zhu, M., Zhang, X., Niu, N. & Shi, Z. METTL14 mediates the m6A methylation of miR-29a-3p, thereby activating the MAP2K6 signaling pathway and exacerbating the inflammatory response associated with spinal tuberculosis. *J*. Orthop. Surg. 20, 625 (2025).

121. Ueda, Y. et al. Interactions of beta-lactam antibiotics and antineoplastic agents. Antimicrob. Agents Chemother. 23, 374–378 (1983).

122. Meyer, F. M. & Bramkamp, M. Cell wall synthesizing complexes in Mycobacteriales. Curr. Opin. Microbiol. 79, 102478 (2024).

123. Sanapalli, B. K. R., Jones, C. R. & Sanapalli, V. Targeting Bacterial Cell Wall Synthesis: Structural Insights and Emerging Therapeutic Strategies. Pharmaceutics 18, 106 (2026).

124. Kumar, K. et al. Discovery of Anti-TB Agents that Target the Cell-Division Protein FtsZ. Future Med. Chem. 2, 1305–1323 (2010).

125. Kwon, K. W. et al. Host-directed anti-mycobacterial activity of colchicine, an anti-gout drug, via strengthened host innate resistance reinforced by the IL-1β/PGE_2_ axis. Br. J. Pharmacol. 179, 3951–3969 (2022).

126. Yao, J. & Rock, C. O. Bacterial fatty acid metabolism in modern antibiotic discovery. Biochim. Biophys. Acta BBA - Mol. Cell Biol. Lipids 1862, 1300–1309 (2017).

127. Gan, Y., Wang, J., Coselli, J. & Wang, X. L. Synergistic induction of apoptosis by HMG-CoA reductase inhibitor and histone deacetylases inhibitor in HeLa cells. Biochem. Biophys. Res. Commun. 365, 386–392 (2008).

128. Heo, S.-K. et al. Targeting c-KIT (CD117) by dasatinib and radotinib promotes acute myeloid leukemia cell death. Sci. Rep. 7, 15278 (2017).

129. Guerrouahen, B. S. et al. Dasatinib Inhibits the Growth of Molecularly Heterogeneous Myeloid Leukemias. Clin. Cancer Res. 16, 1149–1158 (2010).

130. Matsui, K. et al. Stimulation of alpha2-adrenergic receptors impairs influenza virus infection. Sci. Rep. 8, 4631 (2018).

131. Hamilton, J. L. et al. The Association of an Alpha-2 Adrenergic Receptor Agonist and Mortality in Patients With COVID-19. Front. Med. 8, 797647 (2022).

132. Cheng, H. W. The immunomodulatory effects of clonidine, an α-2-adrenergic agonist, in laying hens. Poult. Sci. 85, 452–456 (2006).

133. Jeon, M. et al. Transforming L1000 profiles to RNA-seq-like profiles with deep learning. BMC Bioinformatics 23, 374 (2022).

134. Belloc, F. et al. The stem cell factor–c-KIT pathway must be inhibited to enable apoptosis induced by BCR–ABL inhibitors in chronic myelogenous leukemia cells. Leukemia 23, 679–685 (2009).

135. Wallace, J. et al. Discovery of Bacterial Fatty Acid Synthase Type II Inhibitors Using a Novel Cellular Bioluminescent Reporter Assay. Antimicrob. Agents Chemother. 59, 5775–5787 (2015).

136. Zhang, Y.-M., White, S. W. & Rock, C. O. Inhibiting Bacterial Fatty Acid Synthesis. J. Biol. Chem. 281, 17541–17544 (2006).

