## Supplemental material for "Integrative transcriptome-based drug repurposing in tuberculosis"

[1]Computational Bioscience Program, School of Medicine, University of Colorado Anschutz Medical Campus, Aurora, CO, USA, [2]Computer Science undergraduate program, Metropolitan State University, Denver, CO, [3]Data Science undergraduate program, Michigan State University, East Lansing, MI, USA, [4]Biomedical Laboratory Science undergraduate program, Michigan State University, East Lansing, MI, USA, [5]Department of Biomedical Informatics, Center for Health Artificial Intelligence, University of Colorado Anschutz Medical Campus, Aurora, CO, USA.

**Supplementary information** **2**

1. Additional materials and methods 2
    - 1.1 Baseline comparison 2
        - 1.1.1 Data collection and preprocessing 2
        - 1.1.2 Baseline similarity assessment 2
2. Supplementary Figures 3
    - Figure S1. Assessment of heterogeneity in the individual TB signatures via gene- and pathway overlaps. 3
    - Figure S2. Signature aggregation scheme. 4
    - Figure S3. Variance across individual signatures by technology. 5
    - Figure S4. Top variable GO:BP terms show no enrichment driven by single factors across TB signatures 6
    - Figure S5. Top 20 GO:BP terms not well captured by the aggregated signatures. 7
    - Figure S6. Evaluation of pathway-level deviation between origin-specific aggregated and individual signatures across technologies. 8
    - Figure S7. Cholesterol- and cytokine-related disease-drug pathway networks. 11
    - Figure S8. Frequency of mechanisms of action (MOAs) associated with our 64 high-confidence candidates. 12
    - Figure S9. Pearson correlation of baseline drug-drug and disease-drug samples by tissue types. 14
3. Supplementary Tables 15
    - Table S1. List of public TB datasets used in this work. 15
    - Table S2. Full list of the top 64 high-confidence candidates with score-level support across microarray and RNAseq signatures. 17
    - Table S3. Key differentiating upregulated GO biological process terms enriched in circulating-other comparison 19
    - Table S4. Key differentiating GO biological process terms enriched in primary-cell-line comparison 19
    - Table S5. Dominant GO:BP terms exhibit statistically lower deviation across technologies 20
    - Table S6. Cell line-to-tissue mapping for LINCS drug control samples used in baseline comparisons. 21
4. Supplementary references 22

### Supplementary information

#### 1. Additional materials and methods

##### 1.1 Baseline comparison

To best determine the most biologically compatible pairs of expression baselines for disease-drug signature comparison, we assessed similarity between each disease and drug control sample using Pearson, Spearman[1], Rank Biased Overlap (RBO)[2], and least absolute shrinkage and selection operator (LASSO)[3] approaches.

###### 1.1.1 Data collection and preprocessing

The drug control samples, i.e., gene expression of untreated cell lines, were obtained from LINCS level 3 GSE92742[4]. We excluded expression of non-landmark genes; therefore, we included only the 978 landmark genes for the analyses. We corrected the drug data distribution by performing a quantile normalization on all the drug control samples. Then, a 'target drug profile' was randomly sampled from the normalized drug data and used as the reference vector for the control disease data distribution mapping using quantile transformation; separately applied on the microarray and RNAseq data.

###### 1.1.2 Baseline similarity assessment

Summarized similarity coefficients of each pairwise disease-drug control sample were calculated using the following metrics:

1. Pearson and Spearman correlation

Pearson and Spearman correlation assess how strong the linear relationship between a pair of drug and disease baselines is. While Pearson only considers the overall agreement trend of genes based on their expression values by looking at how well each gene aligns with their respective sample mean, regardless of direction, Spearman correlation is a directional metric that takes into account the difference in ranks of the same gene from two baseline samples.

2. Rank-Biased Overlap (RBO)

Unlike Pearson and Spearman, which require two complete lists for comparison, RBO is a ranked-based measure with weight assignments to all the genes in each baseline sample. The genes ranked toward the top of the list based on absolute regulation level (highly up- or downregulated) get higher weights, meaning if a gene is ranked the same or very close toward the top, then it would upweight the RBO metric. Overall, RBO is claimed to be suitable for feature selection as RBO coefficients are low for the genes with a large difference in ranks, similar to zeroing out unimportant features. Therefore, the RBO metric contribution only considers genes with a high rank agreement between the two lists.

3. Lasso coefficients

We adapted the concept of *SampleLasso*[5] to quantify baseline similarity using L1-regularized regression. For each disease control sample, we trained a Lasso model to predict its expression profile as a sparse linear combination of the drug control cell line profiles. In modeling terms, the cell line profiles served as features, while each disease sample became a target. The resulting Lasso coefficients, by representing the contribution of each cell line profile to reconstructing the disease profile, were used as our similarity metric.

#### 2. Supplementary Figures

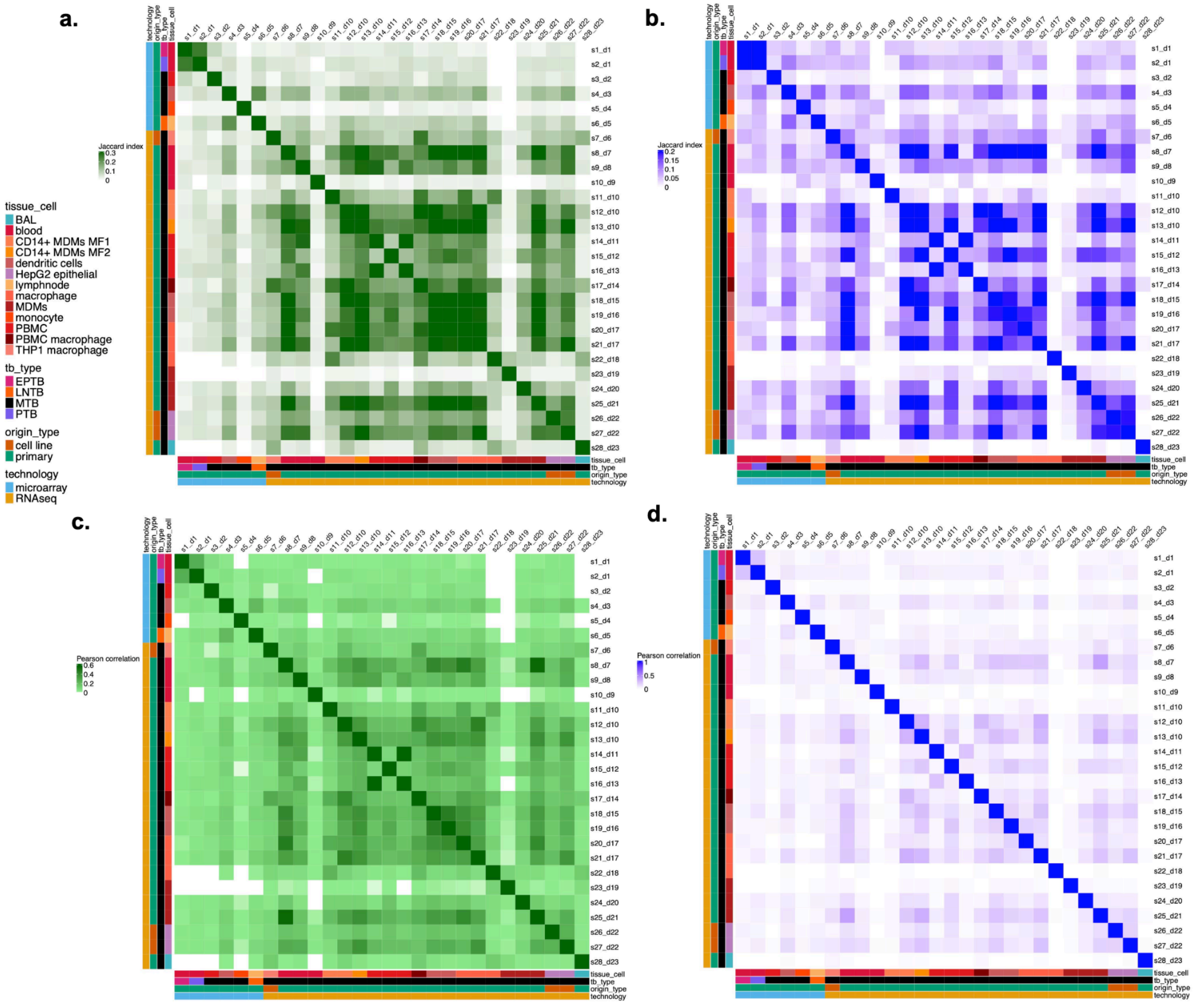


Figure S1. Assessment of heterogeneity in the individual TB signatures via gene- and pathway overlaps.

**(a)** Jaccard similarity matrix of upregulated gene signatures. **(b)** Jaccard similarity matrix of downregulated gene signatures. **(c)** Pearson correlation of asinh-transformed enrichments of upregulated pathway signatures. **(d)** Pearson correlation of asinh-transformed enrichments of downregulated pathway signatures. Details of the signature-dataset identifiers at the tops and right sides of the plots can be found in **Table S1**.

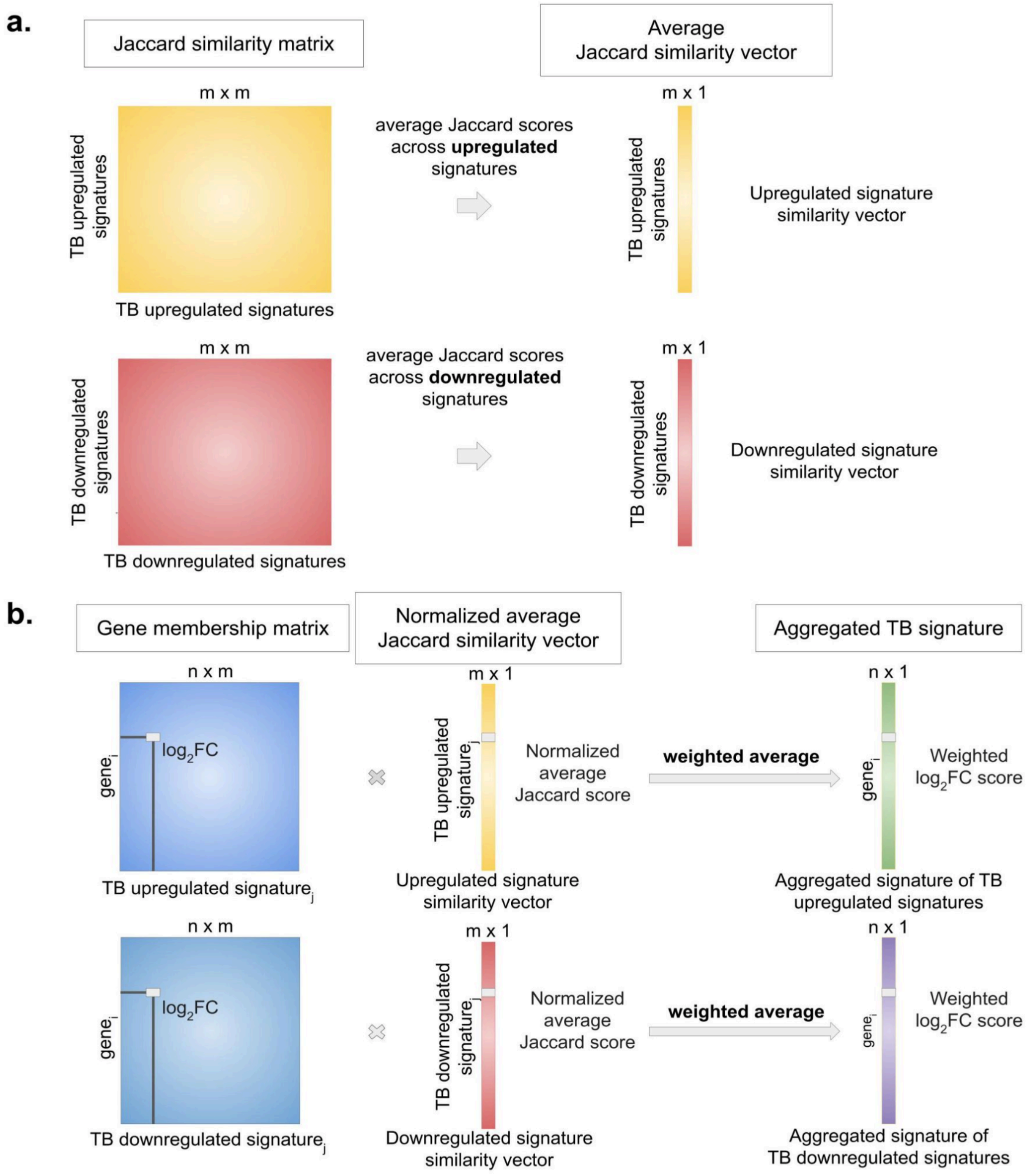


Figure S2. Signature aggregation scheme.

**(a)** Overview of the similarity-weighting step used to assign weights as proxies of ‘confidence’ to each individual TB signature. Each TB signature consists of the top-ranked upregulated and downregulated genes derived from differential expression analysis. Pairwise Jaccard similarity scores were computed between all signatures separately for upregulated (**top**) and downregulated (**bottom**) gene sets to preserve directionality. The average *Jaccard* score for each signature is then used to create a similarity vector representing its agreement with the rest of the group. Signatures with higher average similarity were considered more reproducible and were assigned higher weights, which were subsequently used to scale their contribution in signature aggregation. **(b)** Construction of the aggregated TB signature. Each gene’s $log_2$ fold change ($log_2FC$; differential expression) across individual signatures is combined using a weighted average, where weights are derived from the normalized *Jaccard* similarity vector. This process generates robust aggregated signatures for both up- and down-regulated signature sets that emphasize consistent transcriptomic signals across studies.

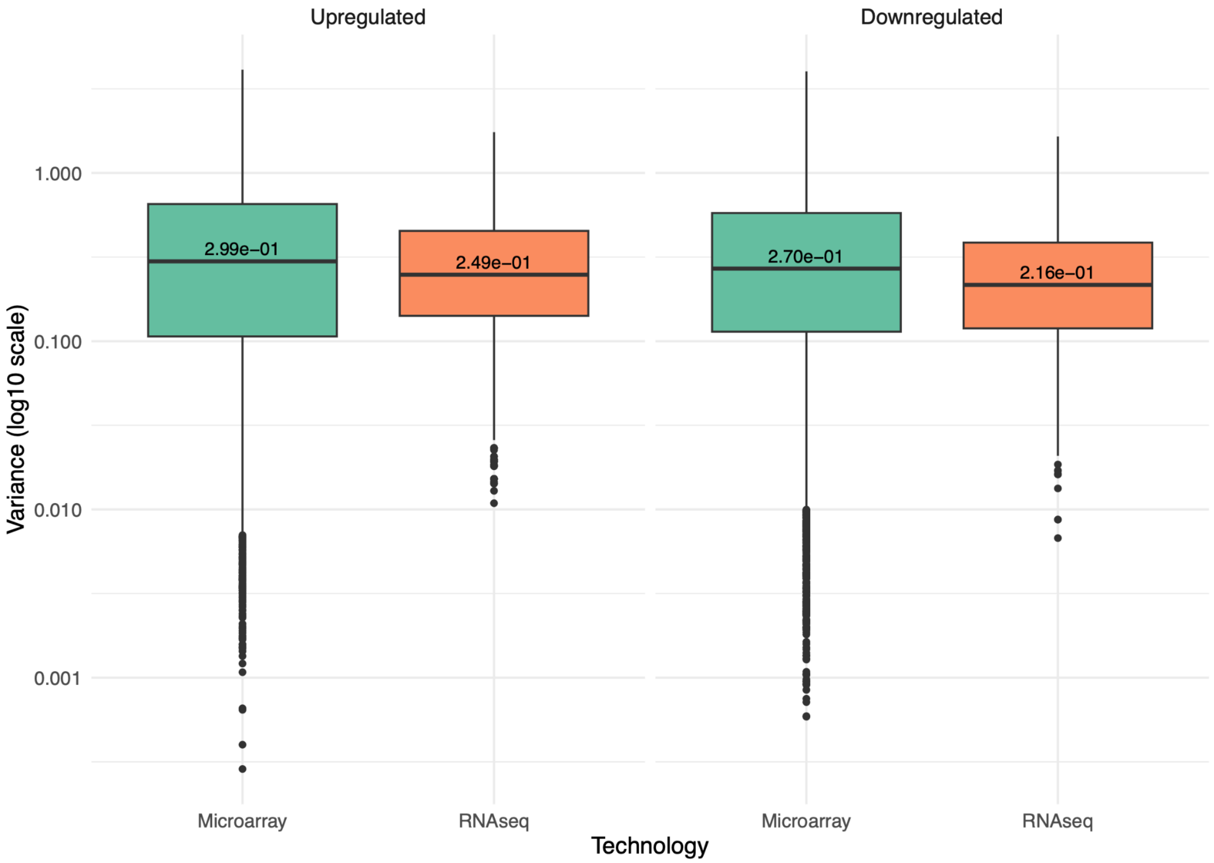


Figure S3. Variance across individual signatures by technology.

Boxplot of log-transformed variance across individual TB signatures derived from 6 microarray (orange) and 22 RNAseq (green) signatures, where each data point represents a pathway. (**left**) Variance distribution of upregulated pathways. (**right**) Variance distribution of downregulated pathways. Higher variance indicates that a pathway was not consistently enriched across individual signatures, reflecting greater heterogeneity in that pathway's involvement across studies.

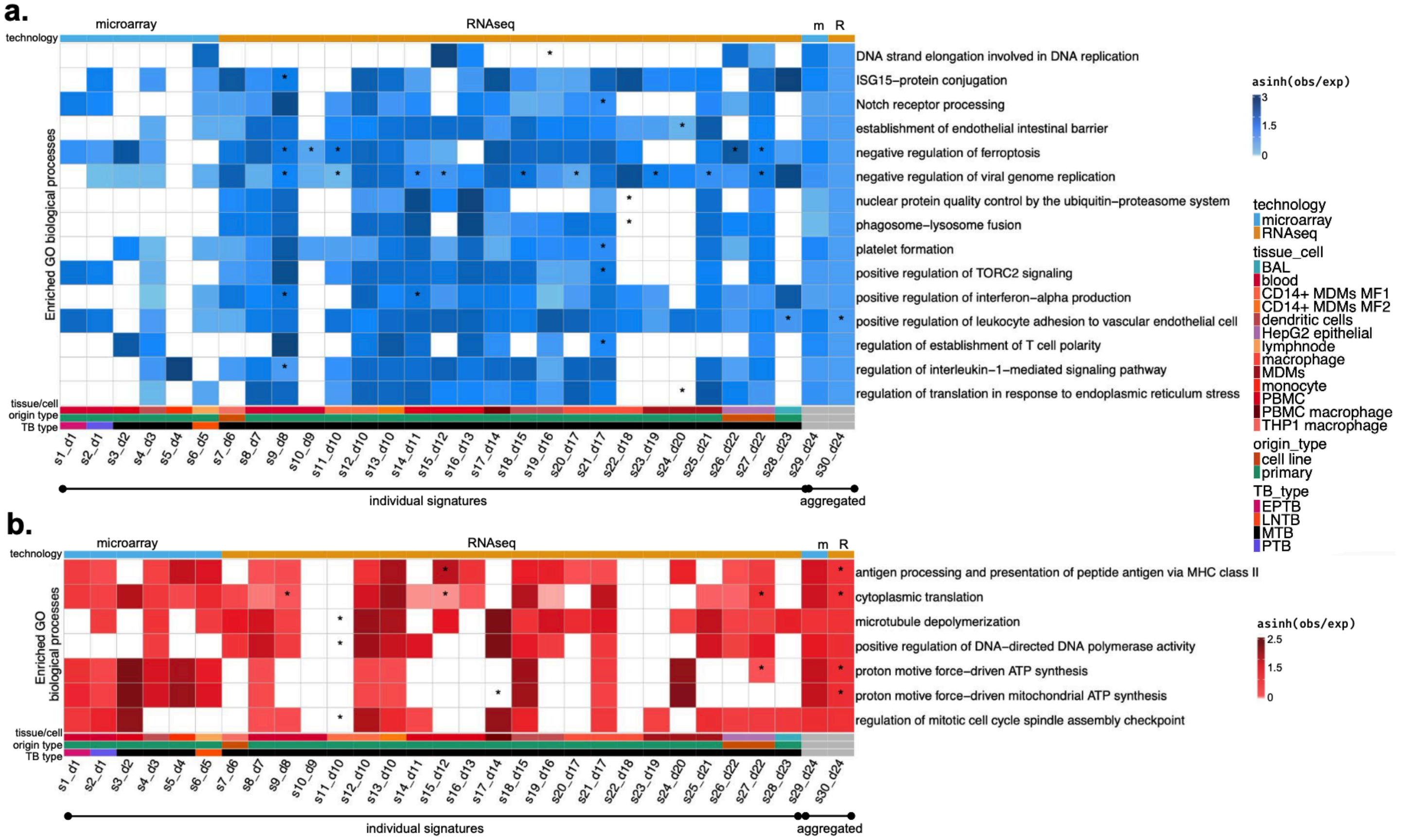


Figure S4. Top variable GO:BP terms show no enrichment driven by single factors across TB signatures

**(a)** Heatmap showing top highly variable upregulated GO biological processes (GO:BP) across the individual TB transcriptomic signatures with asterisks (*) to highlight statistically significant terms (adjusted p-value < 0.05). **(b)** Heatmap showing top highly variable downregulated GO:BP across the individual TB transcriptomic signatures with asterisks to highlight statistically significant terms (adjusted *p-value* < 0.05). The enrichment (i.e., *asinh(observed/expected)*) distributions show that these pathways are inconsistently enriched across signatures. Signature annotations (profiling technology, tissue/cell type, and origin type) are indicated in the annotation bars, highlighting the biological and technical diversity across datasets. Details of the signature-dataset identifiers at the bottom of the plots can be found in **Table S1**.

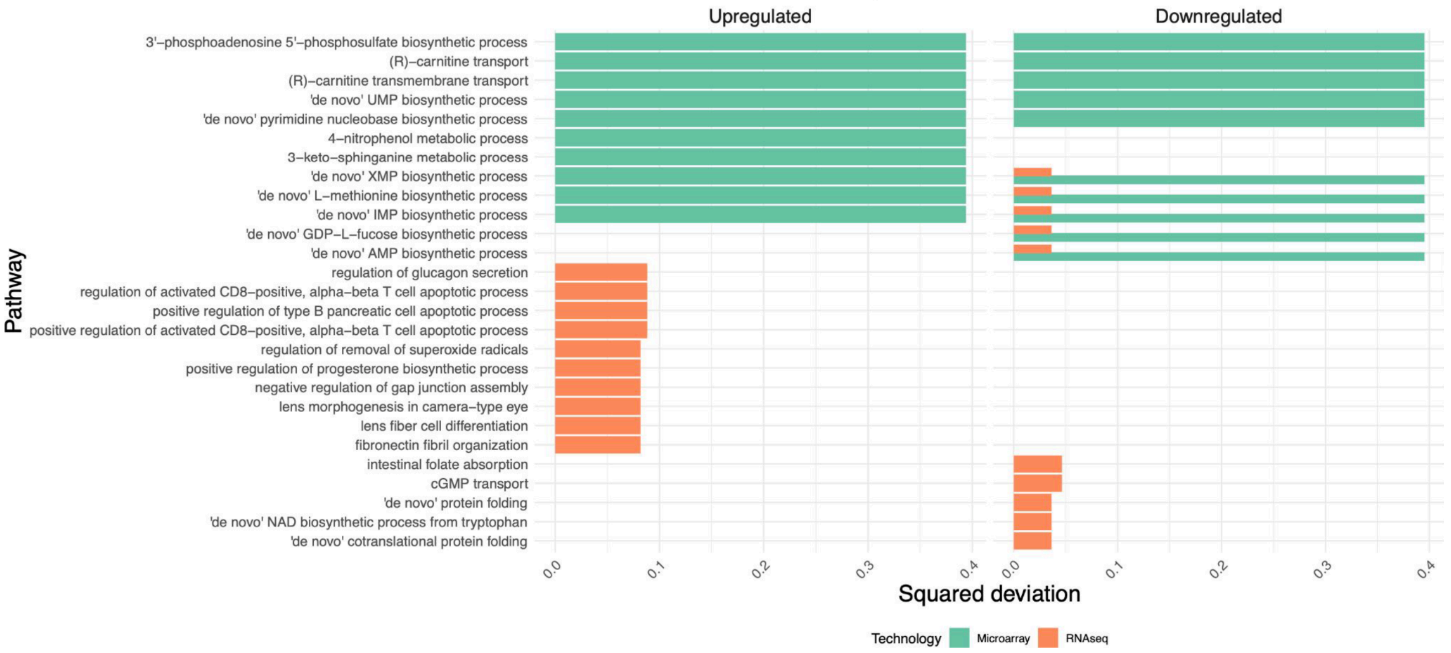


Figure S5. Top 20 GO:BP terms not well captured by the aggregated signatures.

Shared outlier upregulated (**left**) and downregulated (**right**) terms exhibiting the highest squared deviation across both microarray (green) and RNAseq (orange). The x-axis shows the GO:BP terms, and the y-axis represents the scale of squared deviations of enrichment scores of a given term between individual and aggregated signatures.

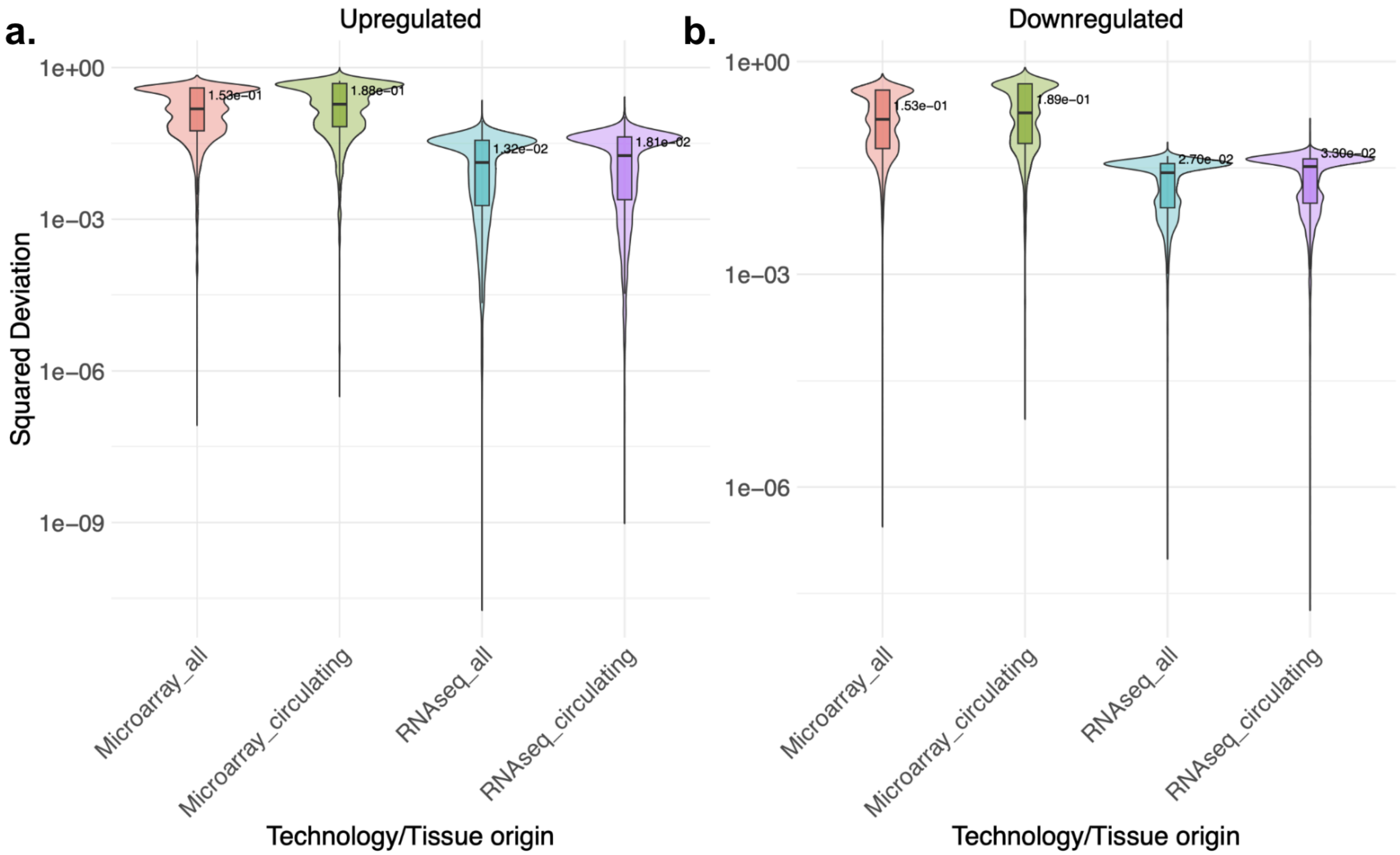


Figure S6. Evaluation of pathway-level deviation between origin-specific aggregated and individual signatures across technologies.

Squared deviation of aggregated pathway enrichment scores from the median of individual signature gene ratios, separately for microarray and RNAseq signatures, separated by regulation direction. (**a**) Upregulated aggregated signatures of all individual signatures vs. circulating-derived individual signatures for microarray (**left**) and RNAseq (**right**). (**b**) Downregulated aggregated signatures of all individual signatures vs. circulating-derived individual signatures for microarray (**left**) and RNAseq (**right**).

a.

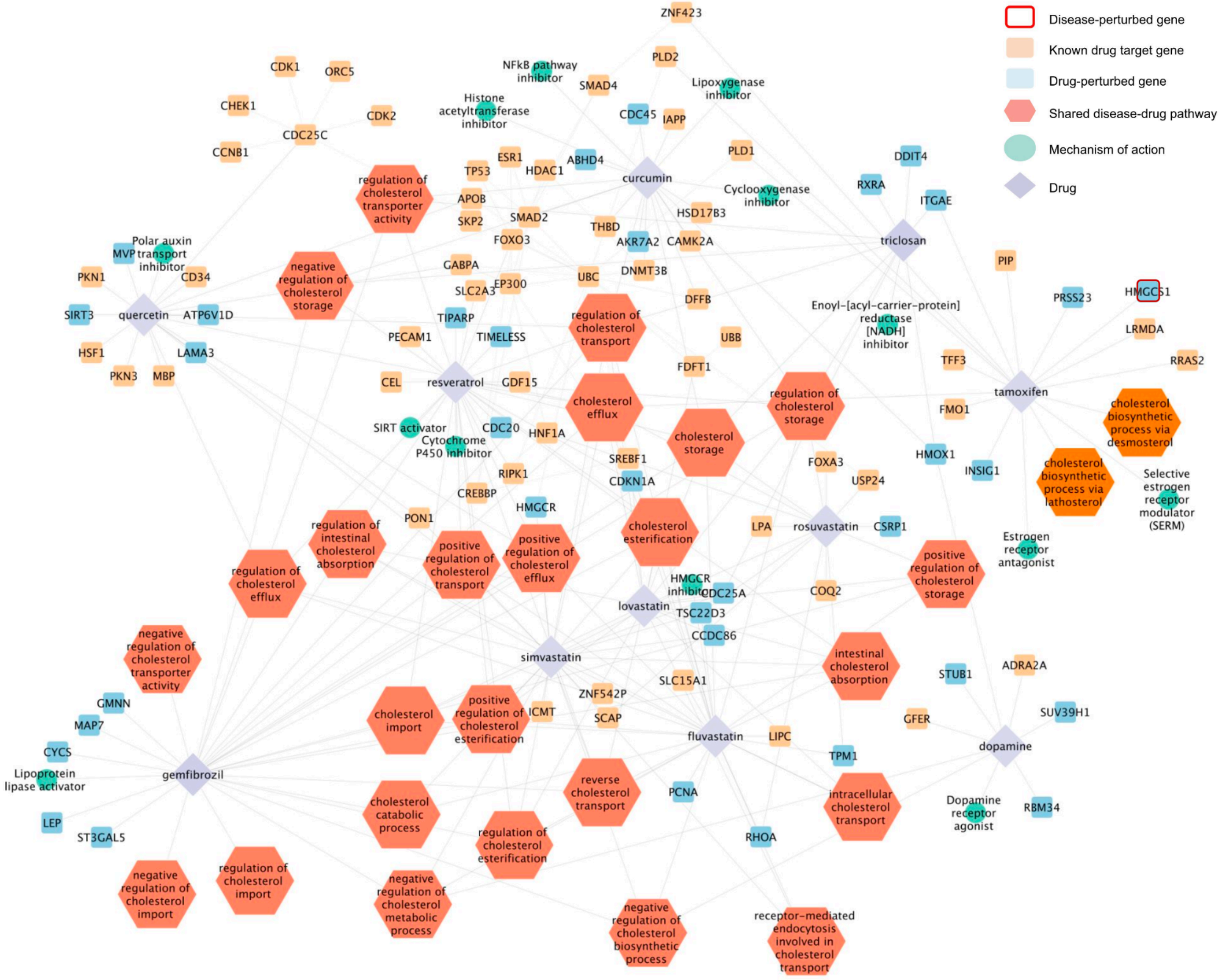

Disease-perturbed gene
Known drug target gene
Drug-perturbed gene
Shared disease-drug pathway
Mechanism of action
Drug
ZNF423
CDK1
ORC5
NFkB pathway inhibitor
PLD2
SMAD4
Lipoxygenase inhibitor
CHEK1
Histone acetyltransferase inhibitor
CDK2
CDC45
IAPP
CDC25C
CCNB1
PLD1
DDIT4
ESR1
ABHD4
TP53
HDAC1
curcumin
regulation of cholesterol transporter activity
RXRA
Cyclooxygenase inhibitor
ITGAE
APOB
SMAD2
HSD17B3
SKP2
THBD
triclosan
FOXO3
AKR7A2
CAMK2A
Polar auxin transport inhibitor
MVP
PKN1
CD34
negative regulation of cholesterol storage
GABPA
PIP
DNMT3B
UBC
EP300
SLC2A3
PRSS23
HMGCS1
DFFB
Enoyl-[acyl-carrier-protein] reductase [NADH] inhibitor
SIRT3
quercetin
ATP6V1D
TIPARP
regulation of cholesterol transport
LRMDA
HSF1
LAMA3
PECAM1
TIMELESS
UBB
RRAS2
PKN3
MBP
FDFT1
TFF3
CEL
resveratrol
GDF15
cholesterol efflux
regulation of cholesterol storage
tamoxifen
cholesterol biosynthetic process via desmosterol
FMO1
SIRT activator
CDC20
HNF1A
Cytochrome P450 inhibitor
cholesterol storage
SREBF1
FOXA3
HMOX1
INSIG1
cholesterol biosynthetic process via lathosterol
Selective estrogen receptor modulator (SERM)
RIPK1
CDKN1A
USP24
CREBBP
HMGCR
PON1
regulation of intestinal cholesterol absorption
cholesterol esterification
LPA
rosuvastatin
CSRP1
Estrogen receptor antagonist
positive regulation of cholesterol transport
positive regulation of cholesterol efflux
regulation of cholesterol efflux
positive regulation of cholesterol storage
HMGCR inhibitor
CDC25A
COQ2
lovastatin
TSC22D3
CCDC86
negative regulation of cholesterol transporter activity
simvastatin
intestinal cholesterol absorption
ADRA2A
STUB1
SLC15A1
ZNF542P
GMNN
positive regulation of cholesterol esterification
ICMT
cholesterol import
SCAP
SUV39H1
MAP7
GFER
fluvastatin
LIPC
dopamine
CYCS
TPM1
Lipoprotein lipase activator
gemfibrozil
PCNA
reverse cholesterol transport
cholesterol catabolic process
intracellular cholesterol transport
Dopamine receptor agonist
RBM34
LEP
ST3GAL5
regulation of cholesterol esterification
RHOA
negative regulation of cholesterol import
regulation of cholesterol import
negative regulation of cholesterol metabolic process
negative regulation of cholesterol biosynthetic process
receptor-mediated endocytosis involved in cholesterol transport

**b.**

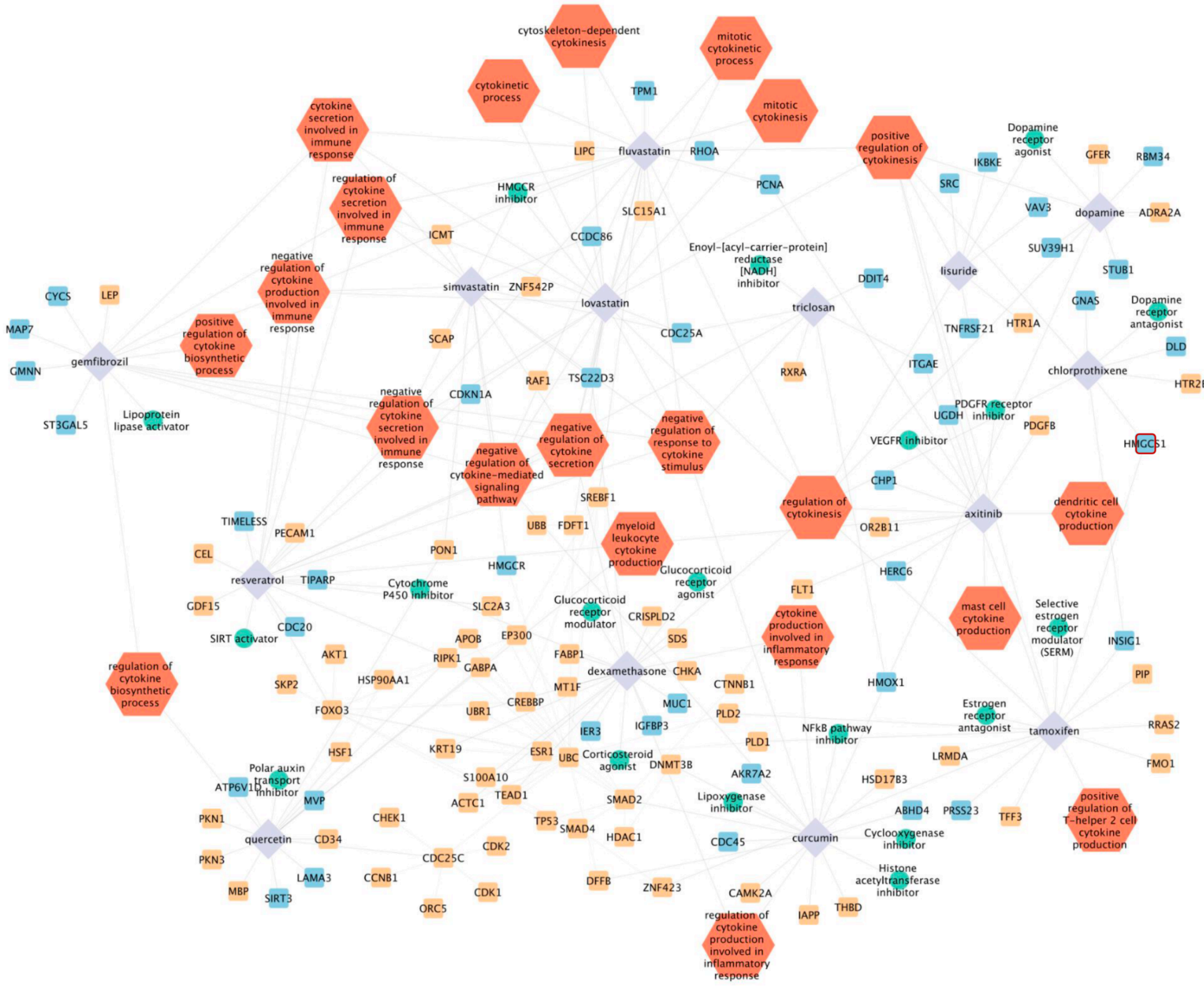
cytoskeleton-dependent cytokinesis
mitotic cytokinetic process
cytokinetic process
TPM1
mitotic cytokinesis
cytokine secretion involved in immune response
LIPC
fluvastatin
RHOA
positive regulation of cytokinesis
Dopamine receptor agonist
GFER
RBM34
IKBKE
regulation of cytokine secretion involved in immune response
HMGCR inhibitor
PCNA
SRC
VAV3
dopamine
ADRA2A
SLC15A1
ICMT
CCDC86
Enoyl-[acyl-carrier-protein] reductase [NADH] inhibitor
SUV39H1
STUB1
negative regulation of cytokine production involved in immune response
DDIT4
lisuride
CYCS
LEP
simvastatin
ZNF542P
lovastatin
triclosan
GNAS
Dopamine receptor antagonist
MAP7
positive regulation of cytokine biosynthetic process
TNFRSF21
HTR1A
SCAP
CDC25A
DLD
GMNN
gemfibrozil
ITGAE
chlorprothixene
RXRA
HTR2B
CDKN1A
RAF1
TSC22D3
ST3GAL5
Lipoprotein lipase activator
negative regulation of cytokine secretion involved in immune response
negative regulation of cytokine secretion
negative regulation of response to cytokine stimulus
UGDH
PDGFR receptor inhibitor
PDGFB
VEGFR inhibitor
HMGCS1
negative regulation of cytokine-mediated signaling pathway
CHP1
SREBF1
regulation of cytokinesis
axitinib
dendritic cell cytokine production
TIMELESS
UBB
FDFT1
myeloid leukocyte cytokine production
OR2B11
PECAM1
CEL
PON1
HMGCR
Glucocorticoid receptor agonist
HERC6
resveratrol
TIPARP
Cytochrome P450 inhibitor
FLT1
GDF15
SLC2A3
Glucocorticoid receptor modulator
CRISPLD2
mast cell cytokine production
Selective estrogen receptor modulator (SERM)
CDC20
cytokine production involved in inflammatory response
SIRT activator
APOB
EP300
SDS
INSIG1
regulation of cytokine biosynthetic process
AKT1
RIPK1
FABP1
GABPA
dexamethasone
CHKA
PIP
SKP2
HSP90AA1
MT1F
CTNNB1
HMOX1
CREBBP
FOXO3
UBR1
MUC1
PLD2
Estrogen receptor antagonist
IER3
IGFBP3
NFkB pathway inhibitor
tamoxifen
RRAS2
HSF1
KRT19
ESR1
UBC
Corticosteroid agonist
PLD1
LRMDA
Polar auxin transport inhibitor
DNMT3B
FMO1
ATP6V1D
S100A10
AKR7A2
HSD17B3
MVP
ACTC1
TEAD1
SMAD2
Lipoxygenase inhibitor
ABHD4
PRSS23
positive regulation of T-helper 2 cell cytokine production
PKN1
CHEK1
TFF3
TP53
SMAD4
quercetin
CD34
HDAC1
CDC45
curcumin
Cyclooxygenase inhibitor
PKN3
CDK2
CDC25C
LAMA3
CCNB1
DFFB
ZNF423
Histone acetyltransferase inhibitor
MBP
SIRT3
CDK1
CAMK2A
ORC5
THBD
IAPP
regulation of cytokine production involved in inflammatory response

Figure S7. Cholesterol- and cytokine-related disease-drug pathway networks.

**(a)** Network centered around cholesterol metabolism, showing interactions among TB disease-perturbed genes in our aggregated disease signatures (red outline), known drug targets (orange), and key shared pathway genes (peach) identified from shortest paths in the STRING protein-protein interaction (STRING-PPI) network. Several drugs and their mechanisms of action (green) converge on cholesterol-related processes, including HMGCR inhibitors and Cytochrome inhibitors. **(b)** Network centered on cytokine-related immune regulation, highlighting shared genes across disease and drug mechanisms, including selective estrogen receptor modulators and tubulin inhibitors.

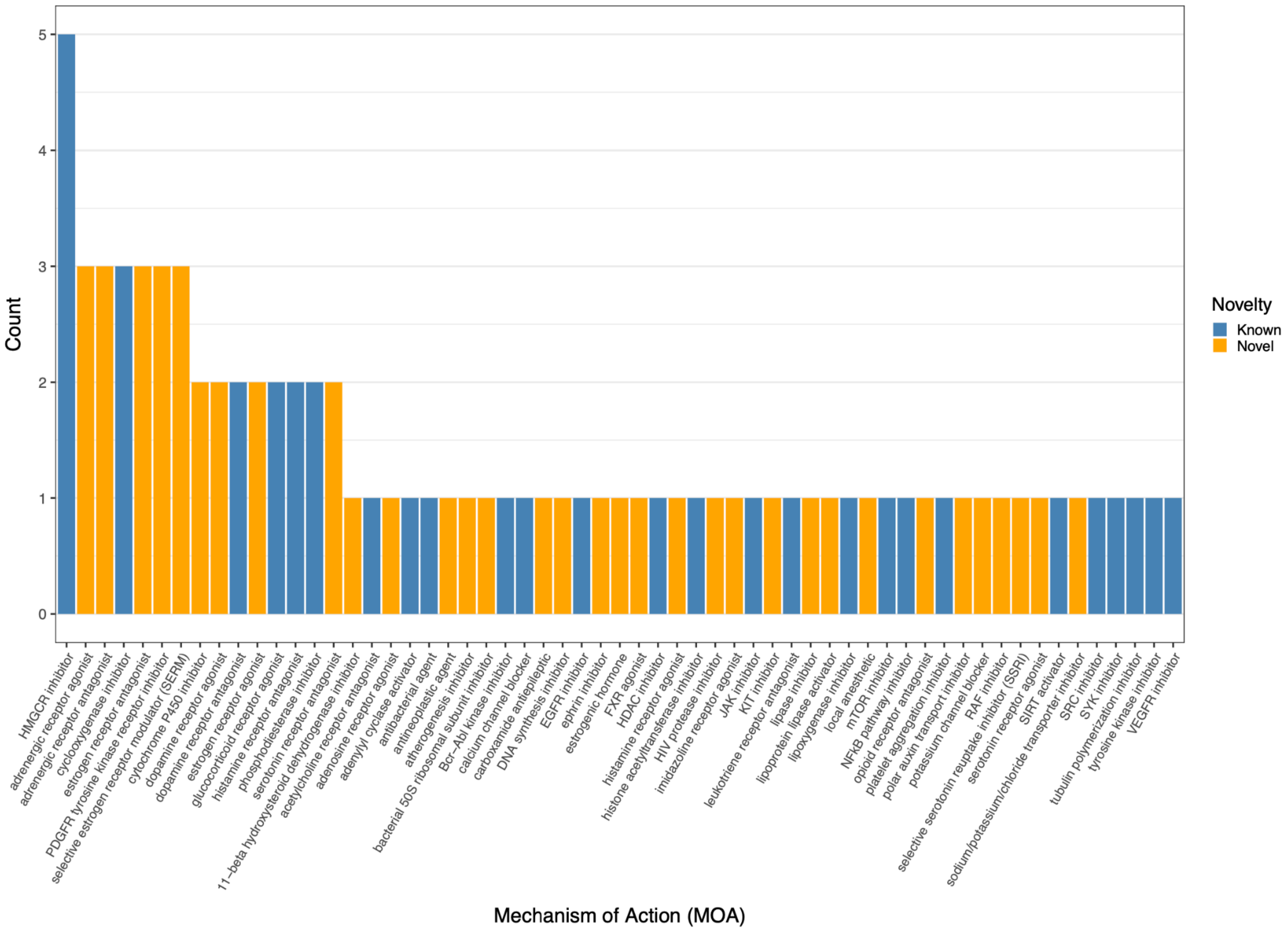


Figure S8. Frequency of mechanisms of action (MOAs) associated with our 64 high-confidence candidates.

Bar plot showing MOAs counts corresponding across the 64 predicted high-confidence drugs coming up in both individual and aggregated signatures in either microarray or RNAseq. The color annotation indicates whether an MOA has been noted as a potential TB HDT by previous research studies (blue: known) or no prior evidence (yellow: novel).

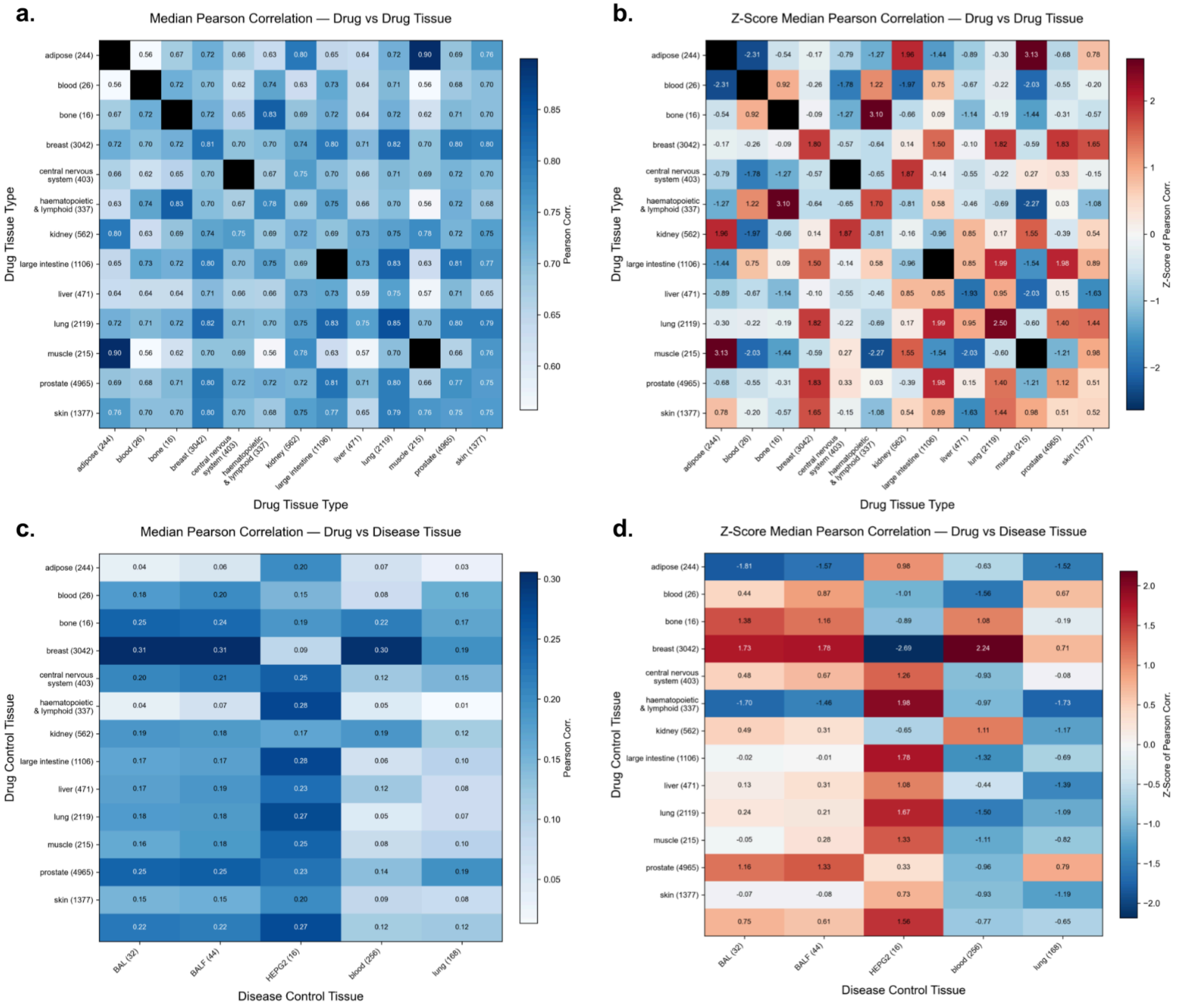


<u>Drug Tissue → Cell Line Mapping</u>

adipose: ASC
blood: U266,
bone: CD34
breast: BT20, HS578T, MCF10A, MCF7, MDAMB231, SKBR3,
central nervous system: NPC
haematopoietic & lymphoid: HL60, JURKAT, NOMO1, THP1, U937
kidney: HA1E, HEK293T, NKDBA
large intestine, HT29
liver: HEPG2, HUH7, PHH
lung: A549, HCC515
muscle: SKB
prostate: PC3, VCAP
skin: A375, FIBRNPC

Figure S9. Pearson correlation of baseline drug-drug and disease-drug samples by tissue types.

**(a)** Heatmaps of raw median Pearson correlations and **(b)** corresponding z-scores between untreated LINCS drug control profiles, grouped by tissue type. Comparisons between identical cell lines were excluded, resulting in null diagonals for tissues represented by a single cell line. Prominent non-null diagonal values reflect the strong similarity among drug control profiles from different cell lines of the same tissue. In addition, the off-diagonal raw values highlight substantial similarity across drug control profiles regardless of tissue type. Many significant off-diagonals were found to be similar tissue type (e.g., carcinoma). **(c)** Heatmaps of raw median Pearson correlations and **(d)** corresponding z-scores between healthy control samples from TB disease datasets (columns) and untreated LINCS drug control profiles (rows), grouped by tissue type. While some tissue-specific patterns are detectable, correlations are consistently lower than those observed among drug control samples, limiting the ability to identify baseline-matched tissue pairs confidently. These results highlight the need for improved approaches to systematically define biologically relevant baselines for disease-drug signature comparisons.

#### 3. Supplementary Tables

##### Table S1. List of public TB datasets used in this work.

This table summarizes the metadata for TB gene expression datasets included in our analysis, grouped by profiling technology (microarray or RNAseq). For each signature, we list the associated study ID, platform, TB status (MTB or PTB), tissue of origin (circulating vs. other), origin type (primary vs. cell line), and cell or tissue type. The number of up- and downregulated genes represents differentially expressed genes used to construct disease signatures. Aggregated signatures represent consensus profiles generated across all microarray or RNAseq studies, respectively.

***Note**: Other studies that did not yield any gene signatures (not included in the table) are microarray: GSE18794, GSE19491, GSE42830, GSE42834, GSE119143, and RNAseq: GSE112482, GSE112483, GSE223863, GSE121049, GSE236156.*

TB signature metadata

Grouped by technology

| No. | Dataset-signature ID | Signature name | TB status | Source | Cell name | Origin type | Tissue origin | Upregulated genes | Downregulated genes | Control samples | Disease samples |
|---|---|---|---|---|---|---|---|---|---|---|---|
| microarray | | | | | | | | | | | |
| 1 | s1_d1 | GSE83456_GPL10558_s1_d1 | PTB | blood | blood | primary | circulating | 54 | 91 | 31 | 45 |
| 2 | s2_d1 | GSE83456_GPL10558_s2_d1 | EPTB | blood | blood | primary | circulating | 44 | 38 | 30 | 47 |
| 3 | s3_d2 | GSE16250_GPL570_s3_d2 | MTB | blood | peripheral blood mononuclear cell | primary | circulating | 56 | 13 | 3 | 3 |
| 4 | s4_d3 | GSE34151_GPL10558_s4_d3 | MTB | blood | dendritic cell | primary | circulating | 179 | 237 | 129 | 130 |
| 5 | s5_d4 | GSE139871_GPL10558_s5_d4 | MTB | blood | monocyte | primary | circulating | 11 | 32 | 4 | 20 |
| 6 | s6_d5 | GSE63548_GPL10558_s6_d5 | LNTB | lymph node | lymph node | primary | other | 108 | 104 | 4 | 22 |
| 29 | s29_d24 | microarray_aggregated_signature | - | - | - | - | - | 98 | 98 | - | - |
| RNAseq | | | | | | | | | | | |
| 7 | s7_d6 | GSE211974_s7_d6 | MTB | THP1 | macrophage | cell line | circulating | 67 | 46 | 3 | 3 |
| 8 | s8_d7 | GSE141656_s8_d7 | MTB | blood | blood | primary | circulating | 267 | 157 | 3 | 3 |
| 9 | s9_d8 | GSE193777_s9_d8 | MTB | blood | blood | primary | circulating | 215 | 108 | 36 | 10 |
| 10 | s10_d9 | GSE84076_s10_d9 | MTB | blood | blood | primary | circulating | 20 | 8 | 4 | 6 |
| 11 | s11_d10 | GSE148731_s11_d10 | MTB | blood | M1 macrophage | primary | circulating | 212 | 147 | 3 | 3 |
| 12 | s12_d10 | GSE148731_s12_d10 | MTB | blood | M1 macrophage | primary | circulating | 50 | 7 | 3 | 3 |
| 13 | s13_d10 | GSE148731_s13_d10 | MTB | blood | M2 macrophage | primary | circulating | 286 | 219 | 4 | 3 |
| 14 | s14_d11 | GSE148171_s14_d11 | MTB | blood | peripheral blood mononuclear cell | primary | circulating | 176 | 52 | 4 | 5 |
| 15 | s15_d12 | GSE174566_s15_d12 | MTxjB | blood | peripheral blood mononuclear cell | primary | circulating | 209 | 95 | 18 | 18 |
| 16 | s16_d13 | GSE198557_s16_d13 | MTB | blood | peripheral blood mononuclear cell | primary | circulating | 156 | 48 | 6 | 6 |
| 17 | s17_d14 | GSE143627_s17_d14 | MTB | blood | mononuclear leukocyte | primary | circulating | 102 | 83 | 3 | 3 |
| 18 | s18_d15 | GSE64179_s18_d15 | MTB | blood | dendritic cell | primary | circulating | 271 | 243 | 6 | 6 |
| 19 | s19_d16 | GSE64182_s19_d16 | MTB | blood | dendritic cell | primary | circulating | 235 | 88 | 3 | 3 |
| 20 | s20_d17 | GSE132283_s20_d17 | MTB | blood | macrophage | primary | circulating | 170 | 54 | 3 | 3 |
| 21 | s21_d17 | GSE132283_s21_d17 | MTB | blood | macrophage | primary | circulating | 336 | 243 | 3 | 3 |
| 22 | s22_d18 | GSE183912_s22_d18 | MTB | blood | macrophage | primary | circulating | 25 | 2 | 4 | 4 |
| 23 | s23_d19 | GSE143731_s23_d19 | MTB | blood | macrophage | primary | circulating | 6 | 7 | 4 | 4 |
| 24 | s24_d20 | GSE164287_s24_d20 | MTB | blood | macrophage | primary | circulating | 81 | 84 | 4 | 4 |
| 25 | s25_d21 | GSE236156_s25_d21 | MTB | blood | macrophage | primary | circulating | 213 | 161 | 11 | 6 |
| 26 | s26_d22 | GSE256184_s26_d22 | MTB | HepG2 | hepatocyte | cell line | other | 410 | 277 | 4 | 4 |
| 27 | s27_d22 | GSE256184_s27_d22 | MTB | HepG2 | hepatocyte | cell line | other | 167 | 124 | 4 | 4 |
| 28 | s28_d23 | GSE165708_s28_d23 | MTB | BALF | alveolar system | primary | other | 23 | 2 | 16 | 16 |
| 30 | s30_d24 | RNAseq_aggregated_signature | - | - | - | - | - | 98 | 98 | - | - |

Table S2. Full list of the top 64 high-confidence candidates with score-level support across microarray and RNAseq signatures.

Each cell shows the number of individual TB signatures (out of 6 for microarray or 22 for RNAseq) for which a given drug achieved a strong reversal score (i.e., drugs ranked in the top 20% most negative minimum reversal scores) under each connectivity metric. The heatmap is split by scoring subcategories: CMAP 1.0, CMAP 2.0 (NCS, Tau, WCS), and correlation-based methods (Pearson, Spearman). Warmer colors (red) represent results from microarray TB signatures; cooler colors (blue) represent RNAseq signatures. Rows correspond to the 64 predicted TB HDT candidates that appeared in both individual and aggregated analyses, and are ranked by their combined mean rank z-score. This visualization highlights which drugs are consistently supported across signatures and metrics, reinforcing their prioritization strength.

| | | | Microarray (6 signatures) | | | | | | RNAseq (22 signatures) | | | | | |
|---|---|---|---|---|---|---|---|---|---|---|---|---|---|---|
| | | | Enrichment-based | | | | Correlation-based | | Enrichment-based | | | | Correlation-based | |
| | | | CMAP1.0 | CMAP2.0 | | | Correlation | | CMAP1.0 | CMAP2.0 | | | Correlation | |
| Rank | combined_z_ | significant_drug | CMAP | NCS | Tau | WCS | XCor | XSpe | CMAP | NCS | Tau | WCS | XCor | XSpe |
| 1 | 3.33 | rosuvastatin | 1 | 1 | 1 | 2 | 0 | 2 | 12 | 12 | 12 | 7 | 9 | 7 |
| 2 | 3.22 | fostamatinib | 3 | 2 | 1 | 0 | 1 | 2 | 18 | 18 | 14 | 5 | 16 | 15 |
| 3 | 2.94 | fluvastatin | 2 | 3 | 2 | 2 | 1 | 2 | 7 | 10 | 10 | 6 | 10 | 7 |
| 4 | 2.85 | tofacitinib | 1 | 0 | 0 | 0 | 1 | 1 | 5 | 9 | 9 | 4 | 5 | 6 |
| 5 | 2.83 | lovastatin | 2 | 2 | 3 | 2 | 2 | 2 | 13 | 12 | 10 | 9 | 7 | 6 |
| 6 | 2.25 | ethinyl-estradiol | 2 | 2 | 2 | 2 | 2 | 0 | 6 | 9 | 7 | 6 | 3 | 3 |
| 7 | 2.18 | atorvastatin | 2 | 3 | 2 | 1 | 2 | 3 | 14 | 10 | 10 | 9 | 10 | 9 |
| 8 | 2.17 | clonidine | 1 | 2 | 1 | 2 | 1 | 1 | 7 | 8 | 7 | 5 | 8 | 6 |
| 9 | 2.16 | nabumetone | 1 | 1 | 2 | 1 | 1 | 1 | 10 | 10 | 6 | 6 | 6 | 5 |
| 10 | 2.05 | tamoxifen | 2 | 2 | 1 | 3 | 2 | 2 | 10 | 11 | 10 | 8 | 11 | 9 |
| 11 | 1.92 | ketotifen | 1 | 2 | 1 | 2 | 2 | 1 | 10 | 9 | 3 | 5 | 6 | 6 |
| 12 | 1.91 | vemurafenib | 2 | 2 | 2 | 1 | 1 | 1 | 17 | 13 | 15 | 6 | 6 | 4 |
| 13 | 1.75 | raloxifene | 1 | 0 | 2 | 0 | 2 | 1 | 13 | 7 | 10 | 6 | 4 | 5 |
| 14 | 1.72 | naltrexone | 2 | 1 | 2 | 1 | 1 | 1 | 11 | 11 | 10 | 3 | 6 | 5 |
| 15 | 1.67 | dasatinib | 1 | 2 | 2 | 2 | 1 | 2 | 12 | 8 | 7 | 4 | 10 | 8 |
| 16 | 1.61 | chlorprothixene | 2 | 2 | 0 | 2 | 0 | 2 | 8 | 11 | 7 | 4 | 4 | 6 |
| 17 | 1.60 | lapatinib | 1 | 2 | 1 | 2 | 2 | 1 | 8 | 8 | 4 | 4 | 4 | 4 |
| 18 | 1.57 | forskolin | 2 | 2 | 2 | 2 | 2 | 3 | 10 | 15 | 14 | 5 | 5 | 9 |
| 19 | 1.53 | sertraline | 2 | 0 | 0 | 1 | 0 | 0 | 17 | 13 | 11 | 5 | 5 | 7 |
| 20 | 1.52 | mometasone | 0 | 2 | 2 | 2 | 1 | 1 | 6 | 6 | 8 | 3 | 7 | 7 |
| 21 | 1.50 | itraconazole | 2 | 2 | 2 | 1 | 2 | 3 | 14 | 12 | 13 | 6 | 7 | 9 |
| 22 | 1.49 | chlorphenamine | 3 | 2 | 1 | 1 | 1 | 2 | 4 | 11 | 11 | 4 | 6 | 5 |
| 23 | 1.45 | NECA | 1 | 2 | 2 | 1 | 1 | 1 | 4 | 10 | 9 | 3 | 4 | 3 |
| 24 | 1.44 | albendazole | 0 | 2 | 2 | 1 | 0 | 2 | 6 | 8 | 8 | 6 | 8 | 7 |
| 25 | 1.44 | quercetin | 0 | 1 | 1 | 1 | 3 | 2 | 9 | 9 | 8 | 7 | 7 | 6 |
| 26 | 1.40 | clarithromycin | 0 | 0 | 1 | 1 | 0 | 0 | 0 | 5 | 11 | 5 | 3 | 4 |
| 27 | 1.33 | temsirolimus | 2 | 1 | 1 | 1 | 1 | 2 | 16 | 13 | 13 | 7 | 2 | 3 |
| 28 | 1.33 | zolmitriptan | 0 | 0 | 2 | 2 | 2 | 1 | 4 | 8 | 4 | 5 | 4 | 6 |
| 29 | 1.33 | panobinostat | 2 | 5 | 4 | 3 | 3 | 2 | 16 | 16 | 14 | 4 | 6 | 8 |
| 30 | 1.32 | resveratrol | 0 | 0 | 1 | 1 | 3 | 2 | 8 | 12 | 13 | 6 | 7 | 9 |
| 31 | 1.26 | nicergoline | 1 | 2 | 2 | 3 | 2 | 2 | 1 | 5 | 5 | 5 | 6 | 7 |
| 32 | 1.14 | icosapent | 1 | 1 | 1 | 1 | 1 | 2 | 8 | 7 | 3 | 4 | 5 | 6 |
| 33 | 1.13 | brompheniramine | 0 | 1 | 0 | 1 | 1 | 1 | 14 | 10 | 11 | 5 | 5 | 6 |
| 34 | 1.10 | dopamine | 1 | 1 | 1 | 1 | 1 | 1 | 2 | 10 | 12 | 5 | 7 | 5 |
| 35 | 1.08 | benzonatate | 2 | 1 | 0 | 1 | 1 | 1 | 8 | 6 | 3 | 5 | 6 | 7 |
| 36 | 1.06 | estradiol | 1 | 2 | 3 | 1 | 2 | 1 | 7 | 5 | 10 | 5 | 7 | 5 |
| 37 | 1.05 | diclofenac | 1 | 1 | 0 | 1 | 2 | 2 | 11 | 6 | 1 | 6 | 7 | 6 |
| 38 | 1.05 | papaverine | 1 | 2 | 1 | 1 | 1 | 1 | 5 | 11 | 10 | 5 | 4 | 7 |
| 39 | 0.98 | oxybenzone | 0 | 0 | 1 | 1 | 2 | 1 | 1 | 3 | 5 | 2 | 5 | 7 |
| 40 | 0.93 | carbamazepine | 0 | 1 | 1 | 0 | 0 | 1 | 4 | 7 | 7 | 4 | 6 | 6 |
| 41 | 0.93 | methoxsalen | 2 | 0 | 0 | 1 | 0 | 1 | 6 | 6 | 5 | 3 | 7 | 3 |
| 42 | 0.91 | ephedrine | 1 | 1 | 1 | 1 | 1 | 1 | 7 | 9 | 12 | 4 | 6 | 8 |
| 43 | 0.91 | curcumin | 0 | 0 | 1 | 0 | 3 | 3 | 7 | 6 | 7 | 4 | 7 | 5 |
| 44 | 0.86 | methylergometrine | 1 | 1 | 1 | 1 | 0 | 2 | 2 | 2 | 3 | 4 | 6 | 5 |
| 45 | 0.84 | tubocurarine | 0 | 1 | 1 | 2 | 2 | 3 | 3 | 6 | 1 | 6 | 5 | 5 |
| 46 | 0.82 | toremifene | 1 | 1 | 1 | 1 | 1 | 2 | 10 | 9 | 7 | 5 | 7 | 8 |
| 47 | 0.79 | naftopidil | 1 | 2 | 3 | 2 | 2 | 1 | 3 | 4 | 5 | 3 | 3 | 6 |
| 48 | 0.67 | probucol | 1 | 1 | 1 | 1 | 2 | 1 | 4 | 6 | 3 | 5 | 4 | 4 |
| 49 | 0.67 | labetalol | 3 | 1 | 0 | 2 | 1 | 2 | 3 | 1 | 1 | 4 | 8 | 5 |
| 50 | 0.59 | simvastatin | 2 | 1 | 0 | 1 | 1 | 2 | 12 | 8 | 6 | 7 | 7 | 8 |
| 51 | 0.53 | bendroflumethiazide | 1 | 0 | 0 | 0 | 1 | 2 | 5 | 7 | 5 | 4 | 6 | 7 |
| 52 | 0.52 | lisuride | 0 | 0 | 0 | 0 | 1 | 1 | 3 | 5 | 8 | 5 | 3 | 4 |
| 53 | 0.51 | nitrendipine | 2 | 3 | 4 | 2 | 2 | 3 | 5 | 9 | 12 | 5 | 6 | 9 |
| 54 | 0.51 | trapidil | 2 | 1 | 1 | 0 | 1 | 1 | 7 | 5 | 5 | 4 | 6 | 5 |
| 55 | 0.49 | dexamethasone | 1 | 1 | 2 | 2 | 2 | 3 | 6 | 6 | 7 | 4 | 6 | 4 |
| 56 | 0.45 | mitotane | 1 | 1 | 0 | 2 | 2 | 2 | 3 | 3 | 3 | 4 | 7 | 4 |
| 57 | 0.39 | amiodarone | 1 | 2 | 1 | 2 | 0 | 2 | 8 | 7 | 5 | 3 | 9 | 8 |
| 58 | 0.33 | triclosan | 0 | 0 | 0 | 1 | 2 | 2 | 4 | 7 | 8 | 3 | 6 | 5 |
| 59 | 0.29 | gemfibrozil | 1 | 2 | 2 | 2 | 2 | 1 | 7 | 6 | 3 | 4 | 2 | 4 |
| 60 | 0.29 | axitinib | 2 | 3 | 1 | 2 | 1 | 1 | 3 | 5 | 1 | 2 | 4 | 4 |
| 61 | 0.23 | nelfinavir | 1 | 0 | 0 | 0 | 2 | 2 | 1 | 1 | 3 | 2 | 4 | 5 |
| 62 | 0.21 | rilmenidine | 1 | 2 | 1 | 2 | 2 | 1 | 5 | 6 | 4 | 6 | 7 | 6 |
| 63 | 0.20 | granisetron | 0 | 0 | 0 | 0 | 0 | 0 | 3 | 8 | 0 | 3 | 1 | 2 |
| 64 | 0.13 | chenodeoxycholic-acid | 0 | 0 | 0 | 0 | 2 | 1 | 2 | 5 | 4 | 5 | 5 | 7 |

Table S3. Key differentiating upregulated GO biological process terms enriched in circulating-other comparison

Top significant GO biological process terms identified by a Mann-Whitney U test comparing enrichment scores (asinh(observed/expected)) between circulating and other TB signatures. Median differences, p-values, and adjusted p-values are reported in scientific notation, rounded to two decimals.

| Top upregulated GO terms enriched in circulating or other | | | | |
|---|---|---|---|---|
| GO term | Enriched in | Median difference (circulating - other) | Raw p-value | Adjusted p-value |
| negative regulation of protection from non-homologous end joining at telomere | Other | −1.47 | $1.43 \times 10^{-5}$ | $2.15 \times 10^{-2}$ |
| negative regulation of telomere maintenance in response to DNA damage | Other | −1.47 | $1.43 \times 10^{-5}$ | $2.15 \times 10^{-2}$ |
| regulation of protection from non-homologous end joining at telomere | Other | −1.47 | $1.43 \times 10^{-5}$ | $2.15 \times 10^{-2}$ |
| regulation of telomere maintenance in response to DNA damage | Other | −1.47 | $1.43 \times 10^{-5}$ | $2.15 \times 10^{-2}$ |
| telomeric DNA-containing double minutes formation | Other | −1.47 | $1.43 \times 10^{-5}$ | $2.15 \times 10^{-2}$ |
| short-chain fatty acid catabolic process | Other | −1.21 | $1.43 \times 10^{-5}$ | $2.15 \times 10^{-2}$ |
| hydrocarbon catabolic process | Other | −1.13 | $1.43 \times 10^{-5}$ | $2.15 \times 10^{-2}$ |

Table S4. Key differentiating GO biological process terms enriched in primary-cell-line comparison

Top significant GO biological process terms identified by a Mann-Whitney U test comparing enrichment scores (asinh(observed/expected)) between primary and cell-line TB signatures. Median differences, p-values, and adjusted p-values are reported in scientific notation, rounded to two decimals.

(a) Upregulated terms

| Top upregulated GO terms enriched in primary or cell line signatures | | | | |
|---|---|---|---|---|
| GO term | Enriched in | Median difference (primary - cell line) | Raw p-value | Adjusted p-value |
| eating behavior | Cell line | −2.40 | $1.12 \times 10^{-5}$ | $1.48 \times 10^{-2}$ |
| reduction of food intake in response to dietary excess | Cell line | −2.40 | $1.12 \times 10^{-5}$ | $1.48 \times 10^{-2}$ |
| response to metformin | Cell line | −2.40 | $1.12 \times 10^{-5}$ | $1.48 \times 10^{-2}$ |
| L-threonine catabolic process to glycine | Cell line | −1.84 | $4.62 \times 10^{-5}$ | $1.48 \times 10^{-2}$ |
| benzoyl-CoA metabolic process | Cell line | −1.84 | $4.62 \times 10^{-5}$ | $1.48 \times 10^{-2}$ |
| establishment of anatomical structure orientation | Cell line | −1.84 | $4.62 \times 10^{-5}$ | $1.48 \times 10^{-2}$ |
| immature B cell differentiation | Cell line | −1.84 | $4.62 \times 10^{-5}$ | $1.48 \times 10^{-2}$ |
| mitochondrial nucleoid organization | Cell line | −1.84 | $4.62 \times 10^{-5}$ | $1.48 \times 10^{-2}$ |
| negative regulation of caveolin-mediated endocytosis | Cell line | −1.84 | $4.62 \times 10^{-5}$ | $1.48 \times 10^{-2}$ |
| negative regulation of cytoplasmic mRNA processing body assembly | Cell line | −1.84 | $4.62 \times 10^{-5}$ | $1.48 \times 10^{-2}$ |

(b) Downregulated terms

| Top downregulated GO terms more enriched in primary or cell line signatures | | | | |
|---|---|---|---|---|
| GO term | Enriched in | Median difference (primary - cell line) | Raw p-value | Adjusted p-value |
| IRES-dependent translational initiation of linear mRNA | Cell line | −1.87 | $4.62 \times 10^{-5}$ | $8.94 \times 10^{-3}$ |
| RNA folding | Cell line | −1.87 | $4.62 \times 10^{-5}$ | $8.94 \times 10^{-3}$ |
| asparagine transmembrane transport | Cell line | −1.87 | $4.62 \times 10^{-5}$ | $8.94 \times 10^{-3}$ |
| cap-independent translational initiation | Cell line | −1.87 | $4.62 \times 10^{-5}$ | $8.94 \times 10^{-3}$ |
| cap-independent translational initiation of linear mRNA | Cell line | −1.87 | $4.62 \times 10^{-5}$ | $8.94 \times 10^{-3}$ |
| cell-cell signaling involved in cell fate commitment | Cell line | −1.87 | $4.62 \times 10^{-5}$ | $8.94 \times 10^{-3}$ |
| chromosome passenger complex localization to spindle midzone | Cell line | −1.87 | $4.62 \times 10^{-5}$ | $8.94 \times 10^{-3}$ |
| exocrine system development | Cell line | −1.87 | $1.29 \times 10^{-4}$ | $2.37 \times 10^{-2}$ |
| extraction of mislocalized protein from ER membrane | Cell line | −1.87 | $4.62 \times 10^{-5}$ | $8.94 \times 10^{-3}$ |
| female sex determination | Cell line | −1.87 | $4.62 \times 10^{-5}$ | $8.94 \times 10^{-3}$ |

#### Table S5. Dominant GO:BP terms exhibit statistically lower deviation across technologies

Mann-Whitney U test results comparing the distributions of deviation scores between dominant (≥90% coverage) and non-dominant (<90% coverage) pathways, stratified by transcriptomic technology and regulation direction. Statistical significance was assessed using Benjamini-Hochberg adjusted p-values, with FDR < 0.05 considered significant.

| Dominant pathways exhibit statistically lower deviation across technologies<br>Mann-Whitney tests comparing ≥90% vs. <90% enriched pathways | | | | | |
|---|---|---|---|---|---|
| Direction | Dominant pathways (≥90% enriched) | Non-dominant pathways (<90% enriched) | Statistic | p-value | Adjusted p-value |
| Microarray | | | | | |
| Upregulated | 1230 | 9941 | $3.23 \times 10^{6}$ | $9.81 \times 10^{-163}$ | $1.31 \times 10^{-162}$ |
| Downregulated | 1050 | 10121 | $2.79 \times 10^{6}$ | $1.58 \times 10^{-143}$ | $1.58 \times 10^{-143}$ |
| RNAseq | | | | | |
| Upregulated | 1798 | 9373 | $2.55 \times 10^{6}$ | 0.00 | 0.00 |
| Downregulated | 518 | 10653 | $8.96 \times 10^{5}$ | $2.79 \times 10^{-163}$ | $5.58 \times 10^{-163}$ |

Table S6. Cell line-to-tissue mapping for LINCS drug control samples used in baseline comparisons.

Each cell line used in the LINCS dataset for untreated drug control profiling was annotated with its corresponding tissue of origin. These mappings were used to assess baseline similarity between disease and drug expression profiles. Cell lines with unknown or ambiguous tissue were removed.

| Cell line | Tissue |
|---|---|
| A375 | skin |
| A549 | lung |
| HCC515 | lung |
| BT20 | breast |
| HME1 | breast |
| HS578T | breast |
| MCF10A | breast |
| MCF 7.00 | breast |
| MDAMB231 | breast |
| SKBR3 | breast |
| HA1E | kidney |
| HELA | large intestine |
| HT29 | large intestine |
| HEPG2 | liver |
| HUVEC | vascular system |
| JURKAT | haematopoietic and lymphoid tissue |
| LNCAP | prostate |
| PC3 | prostate |
| YAPC | pancreas |
| NPC | central nervous system |
| NPC.CAS9 | central nervous system |
| NPC.TAK | central nervous system |
| ASC | adipose |
| ASC.C | adipose |
| CD34 | bone |
| SKL | muscle |
| SKL.C | muscle |

#### 4. Supplementary references


1. Spearman Rank Correlation Coefficient. in *The Concise Encyclopedia of Statistics* 502–505 (Springer New York, New York, NY, 2008). doi:10.1007/978-0-387-32833-1_379.

2. Webber, W., Moffat, A. & Zobel, J. A similarity measure for indefinite rankings. *ACM Trans. Inf. Syst. TOIS* **28**, 1–38 (2010).

3. Tibshirani, R. Regression Shrinkage and Selection Via the Lasso. *J. R. Stat. Soc. Ser. B Stat. Methodol.* **58**, 267–288 (1996).

4. Subramanian, A. *et al.* A Next Generation Connectivity Map: L1000 Platform and the First 1,000,000 Profiles. *Cell* **171**, 1437-1452.e17 (2017).

5. Mancuso, C. A., Canfield, J. L., Singla, D. & Krishnan, A. A flexible, interpretable, and accurate approach for imputing the expression of unmeasured genes. *Nucleic Acids Res.* **48**, e125 (2020).
